# Fine-Tuning of Material Properties via Catch Bonds

**DOI:** 10.1101/2025.01.09.632258

**Authors:** Md Foysal Rabbi, Gijsje H. Koenderink, Yuval Mulla, Taeyoon Kim

## Abstract

Semiflexible polymer networks are ubiquitous in biological systems, including a scaffolding structure within cells called the actin cytoskeleton. The polymers in these networks are often interconnected by transient bonds. For example, actin filaments in the cytoskeleton are physically connected via cross-linkers. The mechanical and kinetic properties of the cross-linkers significantly affect the rheological properties of the actin cytoskeleton. Here, we employed an agent-based model to elucidate how the force-dependent behaviors of the cross-linkers determine the material properties of passive networks without molecular motors and the force generation of active networks with molecular motors. The cross-linkers are assumed to behave as a slip bond whose dissociation rate is proportional to forces or as a catch-slip bond whose dissociation rate is inversely proportional to forces at low force level but proportional to forces at high force level. We found that catch-slip-bond cross-linkers can increase both stress and strain at a yield point by forming force-bearing elements that turn over continuously, which is impossible to achieve without the catch-slip bonds. In addition, we demonstrated that the catch-slip-bond cross-linkers help myosin motors generate greater internal contractile forces by reinforcing the force-bearing parts of the active network.

## INTRODUCTION

Polymeric networks are prevalent in biological systems. In particular, semiflexible networks, consisting of polymers whose persistence length is comparable to their contour length, exist within cells as a scaffolding structure called the cytoskeleton as well as outside cells as extracellular matrices. The mechanical properties of these networks have been extensively investigated in previous studies [1]. Polymers in the semiflexible networks are often interconnected. In the extracellular matrices, polymers may form cross-linking points by themselves [2–4]. Cytoskeletal polymers, such as actin filaments (F-actin) and microtubules, are interconnected mainly via cross-linking proteins that can bind pairs of the cytoskeletal polymers. In the case of the actin cytoskeleton, more than a dozen types of actin cross-linking proteins have been identified [5]. Cross-linking points play an important role in transmitting forces across networks, not only by distributing externally applied forces but also by transmitting internally generated forces to surroundings. For example, in the cortical cytoskeleton located on the inner face of the cell membrane, contractile forces generated by myosin motors are transmitted to cellular scale, mediating various physiological processes including cell migration, cell division, and morphogenesis [6].

Cross-linking points forming between biopolymers are often transient, leading to viscoelastic behaviors of networks, such as stress relaxation in response to applied constant strain [7–9] and creep in response to applied constant stress [10–13]. Most of these viscoelastic responses have been explained by the iterations of the breakage and formation of the cross-linking points. The lifetime of these cross-linking points is mostly force-dependent. For example, the cross-linking points between collagen fibrils break faster if higher forces are applied to the points [12, 14–16], meaning that they behave as a slip bond whose unbinding rate increases with higher forces. While actin cross-linking proteins were generally regarded as the slip bond, some of them, such as α-actinin and filamin A, were found to exhibit more complicated force-dependent unbinding behaviors [17–21]. Their unbinding rate decreases if an applied force increases within a small force range, meaning that they behave as a catch bond like seat belts and Chinese finger traps. The nature of the catch bonds observed in the actin cross-linking proteins originates mainly from their conformational change, such as the exposure of a cryptic binding site. However, their unbinding rate eventually increases if the applied force becomes large enough. The mixed response of slip and catch bonds observed in those cross-linkers is called a catch-slip bond.

Networks can exhibit quite distinct responses depending on whether cross-linking points behave as a slip bond or a catch-slip bond. A previous study showed that polymer-grafted nanoparticle networks with catch bonds exhibit greater ductility, higher toughness, and faster strain recovery than those with slip bonds [22]. Another study showed a difference in the propagation pattern of a crack formed on a network, depending on the nature of the cross-linking points [23]; with slip-bond cross-linking points, a crack kept expanding from an initial location via the subsequent breakage of the cross-linking points occurring mostly at the edges of the crack. On the contrary, a network with catch-slip bonds showed resistance to crack propagation because more cross-linking points were formed at the edges of the crack. In our previous study [18], we demonstrated that an actin network with catch-slip-bond cross-linkers formed a stronger cytoskeletal network whose yield stress and strain are higher than those of a network with slip-bond cross-linkers. We directly adopted the force-dependent unbinding rate of the catch-slip-bond and slip-bond cross-linkers from experimental measurements with wild-type α-actinin, which exhibits the catch-slip behavior, and a K225E point mutant associated with the heritable disease kidney focal segmental glomerulosclerosis type 1, which behaves as the slip bond.

Although our previous study reported the different mechanical behavior of actin networks depending on the nature of cross-linker unbinding, it remains unclear which aspect of the catch-slip bond resulted in higher yield stress and strain. In this study, using a rigorous computational model, we explored various types of catch-slip bonds to illuminate how the mechanical properties of actin networks are fine-tuned by the rearrangement of cross-linkers with the nature of catch-slip bonds. Our model, which can provide critical information at high spatiotemporal level, enabled us to find mechanisms that are impossible to find via only experiments or highly coarse-grained models.

## METHODS

We employed our well-established agent-based model based on Brownian dynamics [24–27]. The details of the model are explained in Supporting Information, and all parameter values are listed in Table S1. Actin filament, cross-linker, and motor are simplified using cylindrical segments (Fig. S1). The filament is simplified into serially connected cylindrical segments with polarity. The cross-linker consists of two segments connected at its center point. The motor consists of a backbone structure with 64 motor arms. Motor backbone segments are fixed in space, whereas segments constituting the filament and the cross-linker change their positions over time. The displacements of the endpoints of the cylindrical segments for the filament and the cross-linker are updated by the Langevin equation and the forward Euler integration scheme. For deterministic forces, three types of forces are considered. Bending and extensional forces maintain angles and distances formed by the cylindrical segments near their equilibrium values, respectively. Repulsive forces account for volume-exclusion effects between overlapping filaments.

A network is created via the dynamic events of actin filaments and cross-linkers, in a three-dimensional rectangular domain (30×30×1 µm) with the periodic boundary condition in all directions. The initial orientations of the filaments are random but perpendicular to the z direction. The cross-linkers bind to pairs of the filaments at a constant rate to form cross-linking points. The cross-linkers also unbind from the filaments in a force-dependent manner; the unbinding rate of the cross-linkers follows either a slip or catch-slip bond behavior (Fig. 1A):

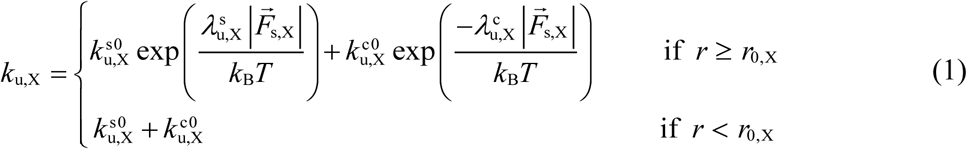

where *k_u,x_^so^* and *k_u,x_^co^* are zero-force unbinding rate constants for slip and catch bonds, respectively, and *λ_u,x_^s^*^s^ and *λ_u,x_^c^* are sensitivity to a tensile force, |*F*_s,x_|, for slip and catch bonds, respectively. In cases considering only slip bonds, *k_u,x_^co^* is set to zero. After unbinding, the cross-linkers undergo the “turnover” event by changing their locations to account for the mobility of the cross-linkers.

**Figure 1.**
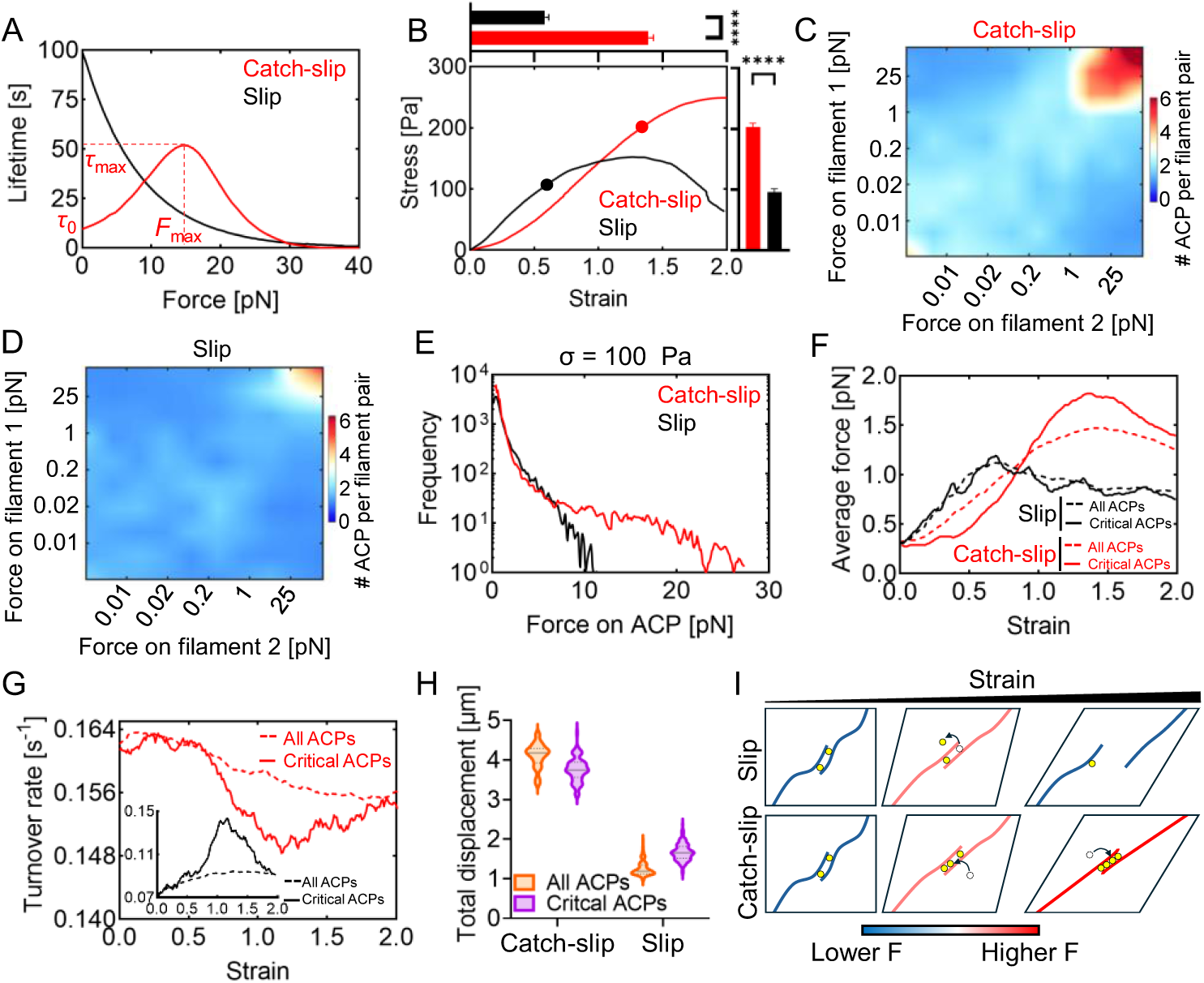
Catch-slip-bond cross-linkers strengthen actin networks via force-dependent redistribution. **A,** Bond lifetime. The lifetime of the catch bond first rises and then decreases with an increasing force, whereas the slip bond shows decreasing lifetime with a higher force. The lifetime of catch-slip bond is characterized by 3 parameters: lifetime with zero force (*τ*_0_), maximal lifetime (*τ*_max_), and force for the maximal lifetime (*F*_max_). Area under curves for these two bonds is identical to each other. **B,** The stress-strain relationship of networks with the catch-slip-bond cross-linkers (red) and the slip-bond cross-linkers (black). Solid circles indicate the yield point in each network. The top panel shows the average yield strain and the right panel shows the average yield stress for each condition. Data are presented as mean values ± s.d. (N = 4 independent samples for each condition). **C-D,** The average number of cross-linkers bound to pairs of filaments bearing different tensile forces. Binning on the x- and y-axes is non-uniform for each bin to contain 10% of filaments. **E,** The distribution of tensile forces on all cross-linkers when network stress was 100 Pa. F-H, A small group of cross-linkers, bearing the top 10% of tensile forces at the yield point, were selected and named critical cross-linkers. The behaviors of critical cross-linkers were compared with those of all cross-linkers. **F,** The average force acting on cross-linkers. In the case with the catch-slip-bond cross-linkers, the average force acting on the critical cross-linkers showed a noticeable deviation from the global average. **G,** The average turnover rate of catch-slip-bond cross-linkers (red). The inset shows the average force acting on slip-bond cross-linkers (black). **H,** The total displacement of cross-linkers. **I,** Different force-dependent redistribution of slip-bond cross-linkers (top) and catch-slip-bond cross-linkers (bottom). Unlike the slip-bond cross-linkers, the catch-slip-bond cross-linkers are redistributed to the part of the network bearing large tensile forces, resulting in larger yield stress and strain. Data are mean ± s.d., n=4. *n* values refer to individual simulation. Statistical analysis was performed using two-sided unpaired *t*-tests **(B)**. ns, not significant; \**P* < 0.05, \*\**P* < 0.01, \*\*\**P* < 0.001, \*\*\*\**P* < 0.0001.

After forming the network, the periodic boundary condition is deactivated in the y direction, and filaments crossing two boundaries normal to the y direction are severed and clamped to the boundaries. Then, two types of studies are performed using the network: bulk rheology and motor contraction. In the bulk rheology study, the network without any motor is subjected to linearly increasing external shear strain as the +y boundary is displaced in the +x direction with the strain rate of 0.001 s^-1^. Shear stress acting on the +y boundary at each strain level is measured. In the motor contraction study, we place a single motor, which mimics the myosin thick filament, at the center of the network and let the motor generate internal contractile forces to the surrounding network. In both types of the measurements, we measure local forces acting on cross-linkers and analyze how the cross-linkers are redistributed across the network.

In most of the simulations, we evaluate a stress-strain relationship via bulk rheology. Unlike linear elastic materials, our networks typically show a non-linear relationship at small strain level with a gradual increase in the slope as soft tissues with fibers [28–30], followed by a linear relationship. Then, at high stress and strain, the slope eventually starts decreasing. The yield point is determined when the slope deviates from the slope of the linear relationship by more than 5%. From the stress-strain relationship, four quantities are calculated: i) initial modulus measured at first 20 s, ii) tangent modulus measured for 10 s before the yield point, iii) stress at the yield point, and iv) strain at the yield point. A network formed by slip-bond cross-linkers is used as a control case. We varied the force-dependent lifetime of catch-slip bonds to understand how the catch-slip bonds tune the material properties of the networks. We also perform simulations with a single motor that generates internal contractile forces in the presence of the slip-bond cross-linkers or the catch-slip-bond cross-linkers to show the generality of our results, regardless of the nature of mechanical loads that networks experience.

## RESULTS AND DISCUSSION

### Catch-slip-bond cross-linkers make stronger actin networks via force-dependent redistribution

First, we performed the bulk rheology measurement using a network formed by slip-bond cross-linkers or catch-slip-bond cross-linkers. The force-dependent lifetime of catch-slip bonds is characterized by 3 parameters: lifetime at zero force (*τ*_0_), maximal lifetime (*τ*_max_), and a force for the maximal lifetime (*F*_max_) (Fig. 1A). Since catch bonds tend to be weaker than slip bonds at low force level [18, 23], we assume that *τ*_0_ of our catch bond is significantly shorter than that of our slip bond. We set the values of these parameters as follows: *τ*_0_ = 9.5 s, *τ*_max_ = 53 s, and *F*_max_ = 14.7 pN. These parameter values enable both catch-slip-bond and slip-bond cross-linkers to have the same total lifetime (i.e., the same area under their force-dependent lifetime curves). There are different sets of the parameter values that can result in the same total lifetime, but we used these specific ones because they lead to the highest *σ*_y_ and *γ*_y_ as we will show later.

We observed significant differences in the stress-strain relationship. In the case with the catch-slip-bond cross-linkers, initial modulus (*E*_0_) was much smaller, whereas tangent modulus measured near the yield point (*E*_y_) was similar (Fig. 1B). Notably, yield stress (*σ*_y_) and yield strain (*γ*_y_) were much larger in the case with the catch-slip-bond cross-linkers (Fig. 1B). As shown in our previous study [18], these differences are involved with how differently cross-linkers are redistributed through the network. When stress reached 100 Pa, we counted the number of the cross-linkers between pairs of filaments as a function of tensile forces that the pairs experience. In the network formed by the catch-slip-bond cross-linkers, there were noticeably more cross-linkers between filaments feeling larger tensile forces than in the network formed by slip-bond cross-linkers (Figs. 1C, D). In addition, in the case of catch-slip-bond cross-linkers, there were more cross-linkers bearing larger tensile forces and fewer cross-linkers supporting smaller tensile forces than those in the case with the slip-bond cross-linkers at the same stress (Fig. 1E). These observations are very similar to those in our previous study [18], and they imply that the catch-slip-bond cross-linkers enable networks to withstand larger stress and strain by reinforcing force-bearing elements in the networks. In this study, we further delved into how the force-dependent cross-linker redistribution takes place.

We identified a small group of cross-linkers bearing the top 10% of the largest tensile forces right before the yield point and compared their properties with those of all cross-linkers. In the case of catch-slip-bond cross-linkers, these critical cross-linkers did not support large forces from the beginning; they experienced smaller forces at lower stress and strain (Fig. 1F). In addition, the average turnover rate of those critical cross-linkers was not noticeably different from the global average turnover rate at small strain but started decreasing as strain increased (Fig. 1G). As a result, the total distance traveled by cross-linker turnover was smaller for the critical cross-linkers (Fig. 1H). Around the yield point, the deviation of the turnover rate of the critical cross-linkers from the global average became maximum, and the average force acting on the critical cross-linkers reached its peak level.

For the slip-bond cross-linkers, we observed quite different tendencies. The average force acting on the critical cross-linkers was similar to the global average force, regardless of strain (Fig. 1F). The turnover rate of cross-linkers increased with higher strain, which is opposite to the observation with the catch-slip-bond cross-linkers (Fig. 1G, inset). The turnover rate of the critical cross-linkers initially followed the global average but then increased faster than the global average as strain increased more. This deviation from the global average peaks near the yield point.

Based on these results, increases in *σ*_y_ and *γ*_y_ induced by catch-slip-bond cross-linkers can be understood as follows (Fig. 1I). Catch-slip bonds show shorter dwell time or more frequent unbinding when they feel no or very small forces. Their unbinding rate becomes minimal, and the dwell time is maximal at intermediate force level. This unique force dependence keeps the catch-slip-bond cross-linkers searching for different regions when stress and strain are low. After they reach regions with intermediate forces at higher stress and strain, their dwell time becomes much longer, resulting in the accumulation of many cross-linkers there. As a result of the accumulation of the catch-slip-bond cross-linkers in high-force regions, the network can hold larger stress and strain before yielding. Slip-bond cross-linkers also undergo the searching process, but in the opposite direction as they tend to stay longer in low-force regions. Thus, the network losing cross-linkers from high-force bearing regions cannot sustain significant stress and strain before yielding.

We repeated simulations using catch-slip-bond cross-linkers with different force dependence (Fig. S2A). The slip-bond part of this new catch-slip-bond cross-linker in Eq. 1 is identical to the equation for slip-bond cross-linkers. This reflects a catch-slip behavior shown by wild-type α-actinin and a slip-bond behavior shown by the K225E point mutant which is linked to an inheritable kidney disease [31]. The force-dependent lifetime of these two cross-linkers is different at low force level but quite similar at high force level, which was assumed in our previous study [18]. New results obtained with the new catch-slip-bond cross-linkers are similar to those shown earlier, meaning that our findings are not dependent on how the force-dependent lifetime of catch-slip bonds is set up between the two options (Fig. S2B-H).

### Network connectivity determines the importance of catch-slip-bond cross-linkers

Here, we probed when increases in *σ*_y_ and *γ*_y_ induced by the redistribution of catch-slip bonds become more effective. We focused on different levels of network connectivity; the network connectivity was varied by changing filament concentration (*R*_F_) or cross-linking density (*R*_X_). It was observed that networks with catch-slip-bond cross-linkers exhibited lower *σ*_y_ and *γ*_y_ than those with slip-bond cross-linkers when the cross-linking density was low (Figs. 2A, B). By contrast, when the cross-linking density was high, networks formed by the catch-slip-bond cross-linkers exhibited greater *σ*_y_ and *γ*_y_ than those with the slip-bond cross-linkers. Interestingly, *σ*_y_ was proportional to both the filament concentration and the cross-linking density, whereas *γ*_y_ showed strong dependence only on the cross-linking density. *γ*_y_ showed an increase only at the low range of the cross-linking density, remaining relatively constant at the high range. We also quantified the turnover rate and force of catch-slip-bond cross-linkers. In networks with high filament concentration, the turnover rate of cross-linkers slowly decreased over time (Fig. 2C, right). The turnover rate of the critical cross-linkers (which bear the top 10% of the largest tensile forces right before the yield point) started deviating from the global average, and the deviation was maximal near the yield point. The critical cross-linkers initially bore smaller forces at lower strain than the global average, reaching peak force level near the yield point (Fig. 2D, right). These overall observations on the turnover rate and the force are similar to those in the case shown earlier (Figs. 1F, G). By contrast, in networks with low filament concentration, the turnover rate increased over time, and the turnover rate of the critical cross-linkers did not show a long-lasting difference from the global average (Fig. 2C, left). In addition, forces acting on the critical cross-linkers were quite similar to the global average (Fig. 2D, left). These tendencies of the turnover rate and the forces share many similarities with those observed with the slip-bond cross-linkers (Figs. 1F and 1G, inset), implying that these cross-linkers failed to settle down in high-force bearing regions. In the following sections, we used the highest filament concentration (10 µM) and the intermediate cross-linking density (0.01) at which the largest long-lasting differences were observed in the turnover and force between two groups of cross-linkers (i.e., the critical cross-linkers vs. all).

**Figure 2.**
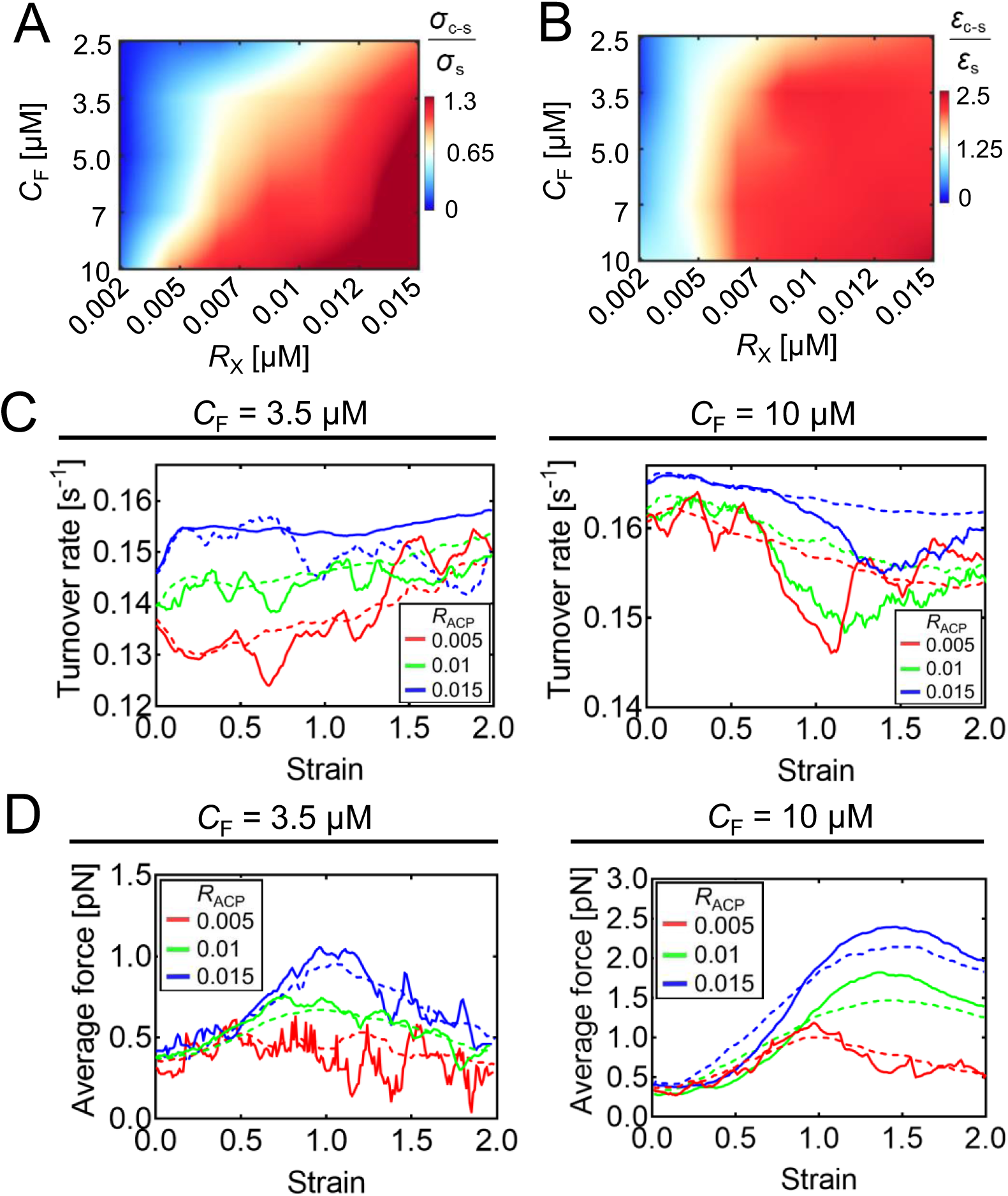
Network strengthening depending on filament concentration (*C*_F_) and cross-linker density (*R*_X_). **A-B,** Comparison of yield stress (*σ*) and yield strain (*ε*) between a network with catch-slip-bond cross-linkers (c-s) and that with the slip-bond cross-linkers (s) with various values of *C*_F_ and *R*_X_. Values greater than one indicate that yield stress or strain in the network with the catch-slip-bond cross-linkers was higher than that in the network with the slip-bond cross-linkers. **C,** The average turnover rate of all cross-linkers (dashed lines) and critical cross-linkers (solid lines), which withstand the top 10% of tensile forces at the yield point, in 6 representative cases. **D,** The average force acting on all cross-linkers (dashed lines) and critical cross-linkers (solid lines) in the same 6 representative cases.

### Catch-slip-bond cross-linkers can increase both stress and strain at the yield point

To better understand how catch-slip-bond cross-linkers increase *σ*_y_ and *γ*_y_, we varied each of the three parameters defining the force-dependent lifetime of the catch-slip bonds from the reference condition (*τ*_0_ = 9.5 s, *τ*_max_ = 53 s, and *F*_max_ = 14.7 pN). First, we changed *F*_max_ with two other parameters (*τ*_0_ and *τ*_max_) fixed and then measured the stress-strain relationship (Figs. 3A, B). We found that the stress-strain relationship at low stress and strain was very similar in terms of magnitude and slope in all cases with different *F*_max_, meaning similar initial network stiffness (= *E*_0_). Network stiffness near the yield point (= *E*_y_) was similar in all cases except one with the lowest *F*_max_ (Fig. 3B & Figs. S3A, B). *σ*_y_ and *γ*_y_ were larger as *F*_max_ was higher (Figs. S3C, D). At the same network-level stress, the distribution of forces acting on all cross-linkers showed a longer tail at high force level with larger *F*_max_ (Fig. S3E). In addition, with higher *F*_max_, the average force acting on critical cross-linkers (which support the top 10% of the largest tensile forces right before the yield point) deviated more from the global average (Fig. 3C); cases with higher *F*_max_ clearly showed much smaller average forces at early times and larger average forces at later times. In the case with the lowest *F*_max_, a difference between the average force of the critical cross-linkers and the global average at small strain level was negligible, implying that the critical cross-linkers did not undergo significant redistribution. In addition, in general, the turnover rate of the cross-linkers was higher with larger *F*_max_ because the range of forces for shorter lifetime on the catch-bond side is wider with larger *F*_max_ (Fig. 3D). With higher *F*_max_, the average turnover rate of the critical cross-linkers began to deviate from the global average from higher strain, but it dropped more substantially before recovering back to the global average. In all cases, the strain level from which the turnover rate of the critical cross-linkers began to drop was close to that where the average force acting on the critical cross-linkers became close to the global average. It is likely that the critical cross-linkers started settling down from this strain level since they began to feel larger forces than the global average.

**Figure 3.**
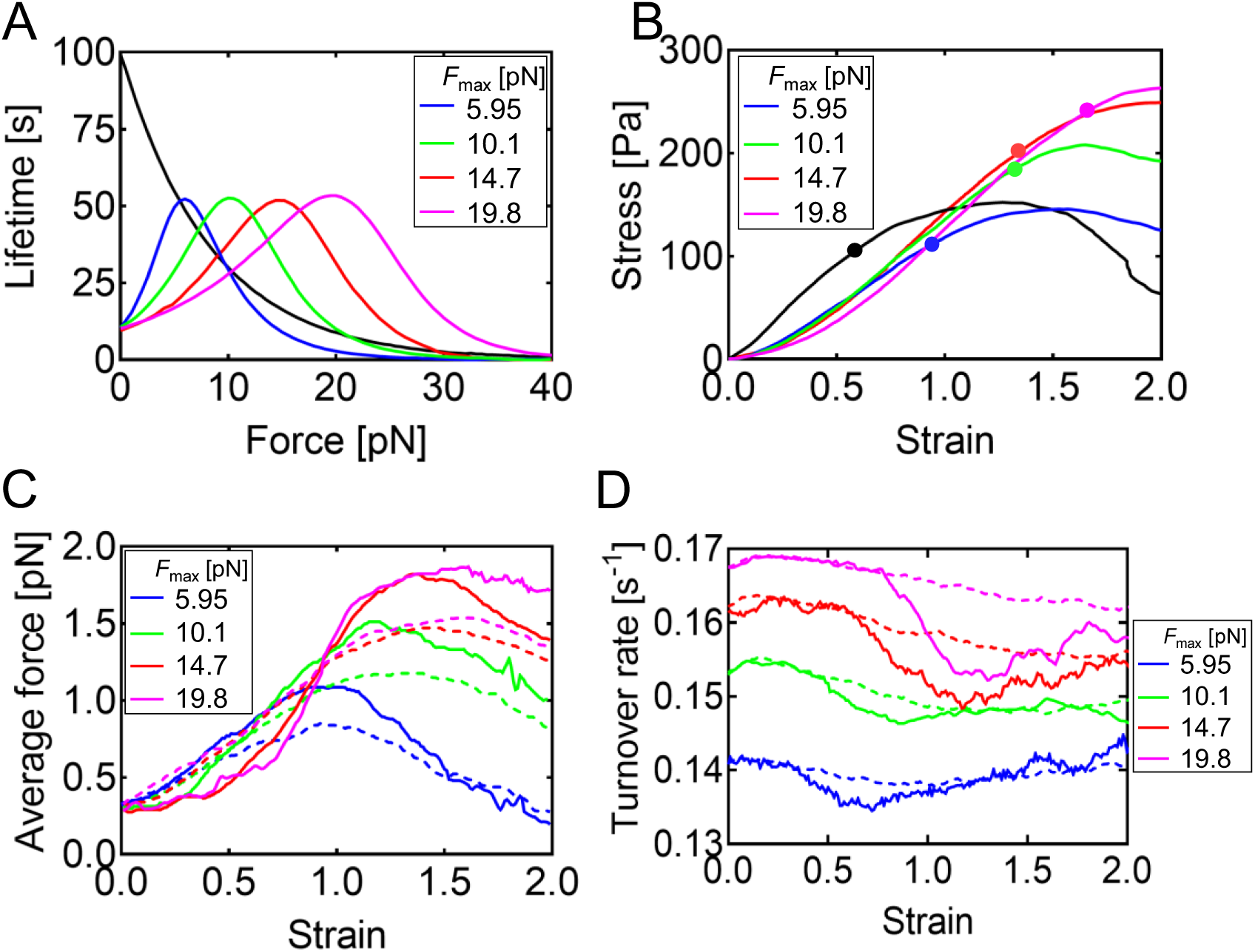
Networks become stronger without a noticeable change in their stiffness if the maximal lifetime of catch-slip bonds emerges at a larger force. **A**, The lifetime of catch-slip bonds with the different force for maximal lifetime (*F*_max_). The black line shows the lifetime of the slip bond as a reference. **B,** The stress-strain relationship with different *F*_max_. The black line shows the stress-strain relationship of the network with slip-bond cross-linkers. Solid circles indicate the yield point in each network. Regardless of *F*_max_, network stiffness represented by a tangent modulus (1/slope) was similar, but yield stress and strain were larger with higher *F*_max_. **C,** The average force acting on all cross-linkers (dashed lines) and critical cross-linkers (solid lines) which withstand the top 10% of tensile forces at the yield point. **D,** The average turnover rate of all cross-linkers (dashed lines) and critical cross-linkers (solid lines). In general, with larger *F*_max_, the average force and turnover rate of the critical cross-linkers showed more deviations from the global averages.

Next, we probed the effect of *τ*_max_ by varying only *τ*_max_ from the reference case (Fig. S4A). Interestingly, *γ*_y_ was similar regardless of *τ*_max_, whereas *σ*_y_ was proportional to *τ*_max_ (Figs. S4B-D). *E*_0_ was similar between the cases, whereas *E*_y_ was apparently larger with higher *τ*_max_. (Figs. S4E, F) Most of the cross-linkers experienced larger forces with higher *τ*_max_ (Fig. S4G). In addition, with higher *τ*_max_, the average force acting on critical cross-linkers deviated more from the global average (Fig. S4H). Except in the case with the lowest *τ*_max_, the average force became similar to the global average at similar strain level, ∼0.9. The turnover rate of the cross-linkers was slightly higher with smaller *τ*_max_ because the lifetime of such bonds is shorter at all force level (Fig. S4I). Regardless of *τ*_max_, the average turnover rate of the critical cross-linkers began to deviate from the global average from similar strain, ∼0.4, but the duration of the deviation was longer with higher *τ*_max_.

Then, we varied *τ*_0_ from the reference condition (Fig. S5A). As *τ*_0_ increased, *γ*_y_ was reduced, whereas *σ*_y_ initially increased and then plateaued (Figs. S5B-D). There was a notable difference in *E*_0_ unlike previous cases with a change in *F*_max_ or *τ*_max_ (Figs. S5B, E, F). With higher *τ*_0_, *E*_0_ was higher, and *E*_y_ was also larger. Crossover between the average force of critical cross-linkers and the global average happened at smaller strain level with higher *τ*_0_ (Fig. S5G). With larger *τ*_0_, the average turnover rate was substantially lower, and the deviation of the average turnover rate of the critical cross-linkers from the global average was smaller (Figs. S5H, I). If *τ*_0_ is high, cross-linkers cannot undergo the force-dependent redistribution process sufficiently because the cross-linkers do not turnover frequently at low force level. Most of the cross-linkers experienced larger forces with higher *τ*_0_ (Fig. S5J).

Among the three parameters, the outcomes of an increase in *F*_max_ are the most interesting. It is not hard to achieve an increase in network stiffness (*E*_0_ or *E*_y_) and an increase in yield stress (*σ*_y_) at the cost of a decrease in strain at the yield point (*γ*_y_), which were observed with an increase in *τ*_max_ or *τ*_0_. For example, even with slip bonds, these changes in the material properties can be achieved simply by increasing the cross-linking density. With more cross-linkers, each cross-linker experiences a smaller load, so a network can sustain larger stress before yielding. The network becomes stiffer because filaments are better connected to each other. However, due to high connectivity, strain at the yield point cannot be large. By contrast, catch-slip-bond cross-linkers with larger *F*_max_ enable a network to have larger stress and strain at the yield point, which cannot be achieved with slip-bond cross-linkers.

### The continuous turnover of force-bearing elements leads to large stress and strain at the yield point

Then, how do cross-linkers with large *F*_max_ enable a network to support large stress and strain at the yield point at the same time? To understand the mechanism, we compared three cases with the largest *F*_max_, *τ*_max_, or *τ*_0_ (Figs. 4A, B). First, we identified filaments bearing top 10% of tensile forces at different strain level in the case with highest *F*_max_. A large fraction of the high-force bearing filaments consistently changed over time, meaning that stress in each strain level was supported by a different set of filaments (Fig. 4C). By contrast, in the case with the highest *τ*_max_, a large portion of the force-bearing filaments were unchanged at higher strain level. In the case with the highest *τ*_0_, the fraction of unchanged force-bearing filaments was nearly 100% at high strain level.

**Figure 4.**
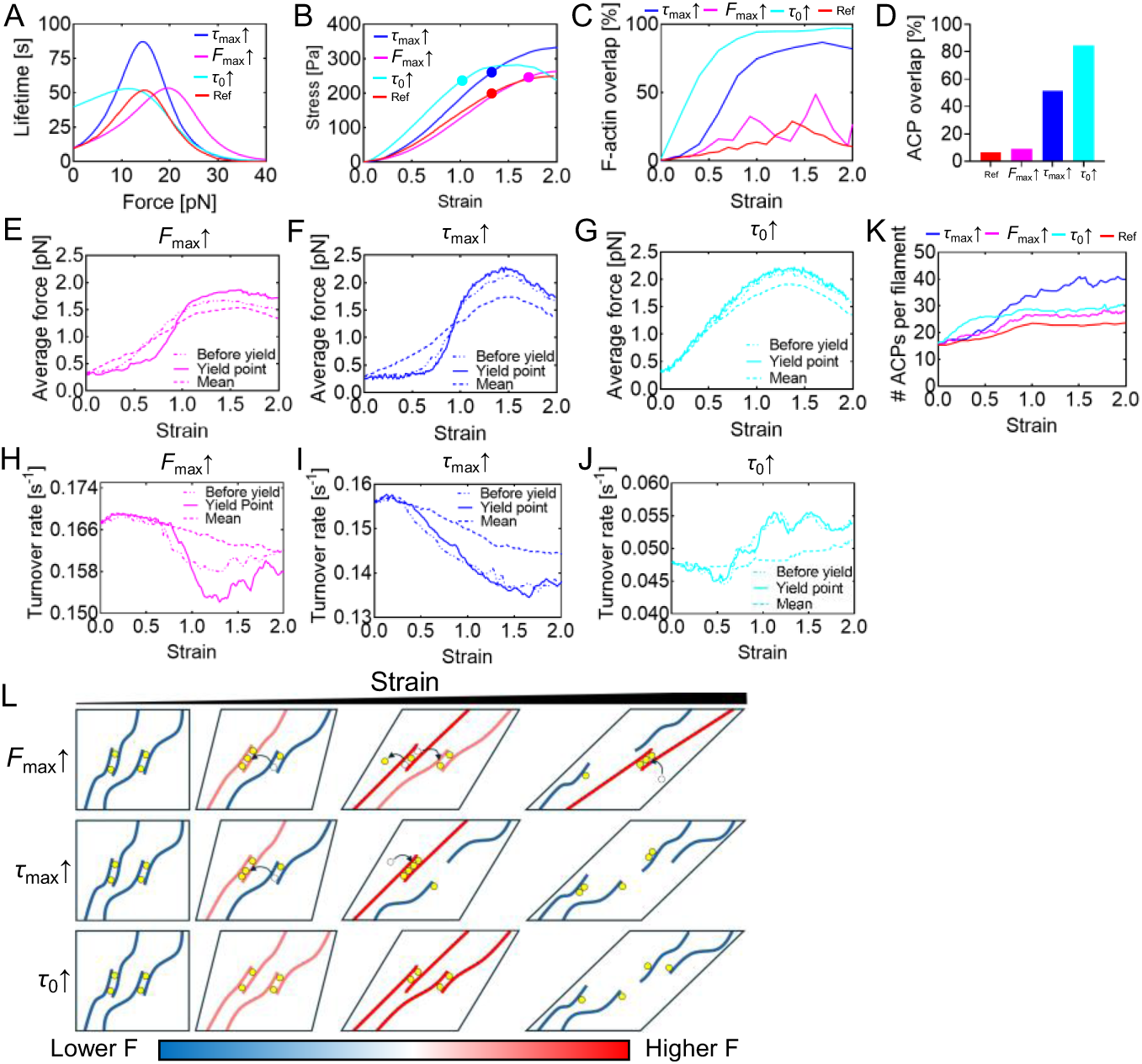
Networks can be strengthened by catch-slip bonds in three different ways. **A,** The case with the largest *F*_max_ (19.8 pN, magenta) is compared to those with the longest *τ*_max_ (88 s, blue) and longest *τ*_0_ (40 s, cyne) to explain the mechanisms. For context, the reference curve is included (*τ*_0_ = 9.5 s, *τ*_max_ = 53 s, and *F*_max_ = 14.7 pN). **B,** The stress-strain relationships for these cases are analyzed and compared. **C,** Percent of overlap between filaments bearing top 10% of tensile forces between two adjacent time points. For example, 100% means that there is no change in filaments in the part of the network bearing high forces. **D,** Percent of overlap between cross-linkers with top 10% of tensile forces selected at the yield point and those identified long before the yield point. From the overlap level of filaments and cross-linkers, it is clear that the high-force bearing part of networks did not change significantly over time in the case with the longest *τ*_max_ and *τ*_0_ unlike the other case. **E-J,** The average force and turnover rate of two groups of cross-linkers identified in D were compared. The former group identified at the yield point is represented by solid lines and “Yield point” in legends, and the latter group is represented by dash-dotted lines and “Before yield” in legends. Global averages are represented by dashed lines and “Mean” in legends. **K,** The average number of cross-linkers bound to filaments bearing top 10% of tensile forces. High-force bearing filaments in the case with the longest *τ*_max_ tended to have more bound cross-linkers. **L,** Three distinct mechanisms of network strengthening by catch-slip bonds. With large *F*_max_, the part of the network bearing large tensile forces keeps changing and has a relatively small number of cross-linkers. The continuous turnover of the high-force bearing part can leads high yield strain. With large *τ*_max_, the high-force bearing part does not change much and keeps recruiting cross-linkers, which results in higher yield stress and relatively the same yield strain. With a highest *τ*_0_, cross-linkers do not undergo force-dependent redistribution at low strain levels. Consequently, the force-bearing elements of the network remain relatively unchanged over time at these lower strain levels. This leads to increased network stiffness from the outset, a behavior not observed in cases with high *F*_max_ and *τ*_max._

In addition, we identified cross-linkers bearing top 10% of the largest tensile forces long before (σ = 30 Pa) the yield point and kept track of the average force acting on them till the yield point. When we compared this group with critical cross-linkers selected at the yield point in the same way, there was only a small overlap between the two groups in the case with the highest *F*_max_ (Fig. 4D), whereas there was a large overlap between the two groups in the cases with the highest *τ*_max_ and *τ*_0._ At the yield point in the case with the highest *F*_max_, cross-linkers in the former group bore lower forces than the critical cross-linkers, implying that the cross-linkers bearing large forces earlier did not support large forces at the yield point anymore (Fig. 4E). By contrast, in the case with the highest *τ*_max_, the two groups of cross-linkers showed similar trends for the average force (Fig. 4F). In the case with the highest *τ*_0_, a significant overlap was present from the beginning, and the average force of the two groups was always equal to or greater than the global average unlike the two other cases (Fig. 4G). The turnover rate of cross-linkers in the two groups also showed similar patterns; in the case with the highest *F*_max_, the turnover rate was different between the two groups, whereas it was very similar between the two groups in the cases with the highest *τ*_max_ and *τ*_0_ (Figs. 4H, I, J).

We also quantified the average number of cross-linkers bound to each of the filaments bearing high forces at each strain level. In the case with the highest *F*_max_, the average number of the cross-linkers bound to high-force bearing filaments did not show a noticeable increase after the strain of ∼1, indicating that there was no continuous accumulation on the same filaments over time (Fig. 4K). On the contrary, the case with the highest *τ*_max_ showed a gradual increase in the average number of the cross-linkers till the end of the simulation, meaning that the cross-linkers kept accumulating on the high-force regions of the network. In the case with the highest *τ*_0_, we observed a fast increase in the average number of cross-linkers per filament at low strain level, which then increased very slowly till the end of the simulation.

As stress increases, forces acting on cross-linkers increase as well. With higher *F*_max_, critical cross-linkers will spend longer time for the searching process via more frequent turnover until they reach forces close to *F*_max_ at higher strain level. Initial network stiffness (*E*_0_) is low because the cross-linkers feeling low forces unbind from filaments frequently. As stress approaches level where critical cross-linkers feel forces close to *F*_max_, they start accumulating near high-force regions with less frequent turnover in order to reinforce the force-bearing parts of the networks near the yield point (Fig. 4L, top). One interesting observation was that networks behave as a material with a relatively constant tangent modulus (*E*_y_), regardless of *F*_max_ (Fig. 3B). This is because the force-bearing elements keep changing over time and maintain a similar number of cross-linkers due to intermediate level of *τ*_max_, rather than lasting for long time with a continuous increase in the number of cross-linkers. This is also crucial for achieving larger *γ*_y_. In our previous study [32], we demonstrated the existence of percolating pathways that support a large portion of forces during network deformation, and the network could show an increase in strain via the continuous turnover of the percolating pathways driven by the turnover of filaments and the unbinding of cross-linkers. Likewise, the time-varying force-bearing elements can lead to a continuous increase in strain. After stress entered higher level which made the critical cross-linkers undergo a transition from the catch phase to the slip phase, they started dissociating from the high-force regions, which eventually resulted in yielding.

How a change in *τ*_max_ or *τ*_0_ affects the material properties is very different from *F*_max_. In networks with higher *τ*_max_, *E*_0_ is still low because high *τ*_max_ does not prevent cross-linkers from undergoing frequent unbinding events at low strain level (Fig. S4B). Once the first force-bearing elements are established, they last longer if *τ*_max_ is higher. Then, additional cross-linkers keep being recruited to the long-lasting force-bearing elements (Fig. 4L, middle). Since the force-bearing elements which constitute percolating pathways hardly changed over time after their first establishment, *γ*_y_ was similar regardless of *τ*_max_. In addition, the existence of more cross-linkers on the force-bearing elements with higher *τ*_max_ resulted in a stiffer network (i.e., higher *E*_y_). In our previous study [26], we have shown the importance of cross-linker stiffness for overall network stiffness. A larger number of cross-linkers on a single cross-linking point are equivalent to a single cross-linker with large stiffness, which explains the larger network stiffness with higher *τ*_max_. After strain reaches higher level which cannot be accommodated by the long-lasting force-bearing elements, high stress makes the cross-linkers in the force-bearing elements undergo a transition from the catch phase to the slip phase, which results in yielding. Because the lifetime of cross-linkers with higher *τ*_max_ is longer in the slip phase, larger stress can be supported until the yield point. With high *τ*_0_, cross-linkers do not undergo the force-dependent redistribution at low strain level. Thus, the force-bearing elements of the networks do not change much over time from the low strain level (Fig. 4L, bottom). This results in high network stiffness from low strain level (i.e., high *E*_0_), which is not observed in the cases with high *F*_max_ and *τ*_max_ (Fig. S5B). Due to percolating pathways with much less frequent turnover, *γ*_y_ becomes very small.

In this study, we fixed the shear strain rate at *γ* = 0.001 s^-1^ which was used in our previous study [18]. It is likely that the rate of. force development on cross-linkers would be proportional to *γ*. If *γ* is larger, force-bearing elements would need to turn over faster to adapt to faster network deformation for larger *γ*_y_. Although changing *γ* is beyond the scope of this study, it is expected that if *γ* becomes higher, *τ*_0_ and *τ*_max_ should be reduced in order to achieve similar *σ*_y_ and *γ*_y_.

### An increase in total bond lifetime or lifetime at high forces does not explain the increase in both stress and strain at the yield point

When we increased *F*_max_ earlier, the total lifetime of the bonds (i.e., the area under the lifetime curve) was also increased significantly. It is possible that the change in the total lifetime also affected the material properties of networks. In particular, the increase in both yield stress and strain might be attributed to an increase in the total lifetime of bonds. To exclude the effect of the total lifetime, we tested five cases with different lifetime profiles with the same total lifetime (Fig. S6A). To maintain the same total lifetime, *τ*_max_ was set to be inversely proportional to *F*_max_, whereas *τ*_0_ was fixed. *E*_y_ was very similar between three cases with lower *F*_max_, but *E*_y_ became lower in two other cases (Fig. S6B). While *γ*_y_ was proportional to *F*_max_, *σ*_y_ showed biphasic dependence on *F*_max_ due to the case with the highest *F*_max_ (Figs. S6C, D). Lower *σ*_y_ and *E*_y_ in those cases with higher *F*_max_ were attributed to low *τ*_max_. We showed earlier that a decrease in *τ*_max_ reduces *E*_y_ and *σ*_y_ (Figs. S4C, F). The reference case with intermediate values of *F*_max_ and *τ*_max_ showed maximal *σ*_y_. This case showed the most distinct deviation of the average force acting on critical cross-linkers and their average turnover rate from the global average values, which explains how the intermediate case could show the highest *σ*_y_ (Figs. S6E, F). In the case with the highest *σ*_y_, the distribution of forces acting on all cross-linkers showed the longest tail (Fig. S6G). From these observations, it is apparent that the change in the total bond lifetime is not a main reason for large stress and strain at the yield point.

When *F*_max_ was increased, bond lifetime at high forces was noticeably increased (Fig. 3A). This change might be a reason for large stress and strain at the yield point rather than the nature of catch-slip bonds. To test whether this is the case or not, we performed simulations with slip bonds with different force dependence of lifetime. While the total lifetime of the slip bonds was fixed, we varied lifetime at zero force over a wide range (Fig. S7A). With lower zero-force lifetime, lifetime at high forces becomes higher. When the zero-force lifetime was 1/2 or 3/4 of the reference value (100 s), both *σ*_y_ and *γ*_y_ were slightly increased, whereas *E*_0_ and *E*_y_ were similar to each other (Fig. S7B). When the zero-force lifetime was 1/4 of the reference value, *γ*_y_ was increased, but *σ*_y_, *E*_0_, and *E*_y_ were reduced. Thus, the longer lifetime at high forces is not a reason for larger stress and strain at the yield point.

These observations imply that the unique force dependence of catch-slip bonds characterized by a difference in lifetime between low and intermediate forces is essential for the increase in both stress and strain at the yield point, rather than higher total bond lifetime or longer lifetime at high force level.

### The force-dependent redistribution of catch-slip bonds takes place and thus enhances force generation in actomyosin networks

So far, we have applied external shear strain to a cross-linked actin network and shown how the force-dependent redistribution of catch-slip-bond cross-linkers can change the material properties of the network. Most types of cross-linked actin networks in cells have myosin motors [33]. These motors generate internal contractile forces to filaments and cross-linkers in the networks. Then, the force-dependent redistribution of the cross-linkers may occur, affecting the force generation process. We probed whether the redistribution of the cross-linkers can take place in an active network with a molecular motor in the absence of external strain. Although our model can include multiple mobile motors in a network as shown in previous works [34–36], we placed a single immobile motor at the center of the network to generate internal contractile forces to the network in a controlled manner. We tested only the catch-slip-bond cross-linkers and the slip bond cross-linkers shown in Fig. 1A. It was observed that a network with the catch-slip-bond cross-linkers generated larger internal stress, and contractile forces propagated over longer distances (Figs. 5A, B), implying connectivity between filaments was better. The catch-slip-bond cross-linkers tended to support much larger forces than the slip-bond cross-linkers (Figs. 5C, D). The average turnover rate and average displacement of critical cross-linkers identified at the yield point using top 10% of tensile forces became lower than the global averages in the case with the catch-slip-bond cross-linkers, whereas they were higher than the global averages in the case with the slip-bond cross-linkers (Figs. 5E, F). We observed more accumulation of the cross-linkers around the motor in the network with the catch-slip-bond cross-linkers (Fig. 5G). In the network with the slip-bond cross-linkers, cross-linker density was higher in locations far from the motor. It is possible that the cross-linkers were accumulated more around the motor due to stronger network contraction induced by larger internal stress, not due to the force-dependent redistribution of the cross-linkers. Thus, we analyzed which density increased earlier between the filaments and the cross-linkers (Figs. 5H, I). In the network with the catch-slip-bond cross-linkers, cross-linker density increased faster than filament density, meaning that those cross-linkers arrived near the motor even before network contraction. By contrast, in the network with the slip-bond cross-linkers, filament accumulation happened earlier than an increase in the cross-linker density.

**Figure 5.**
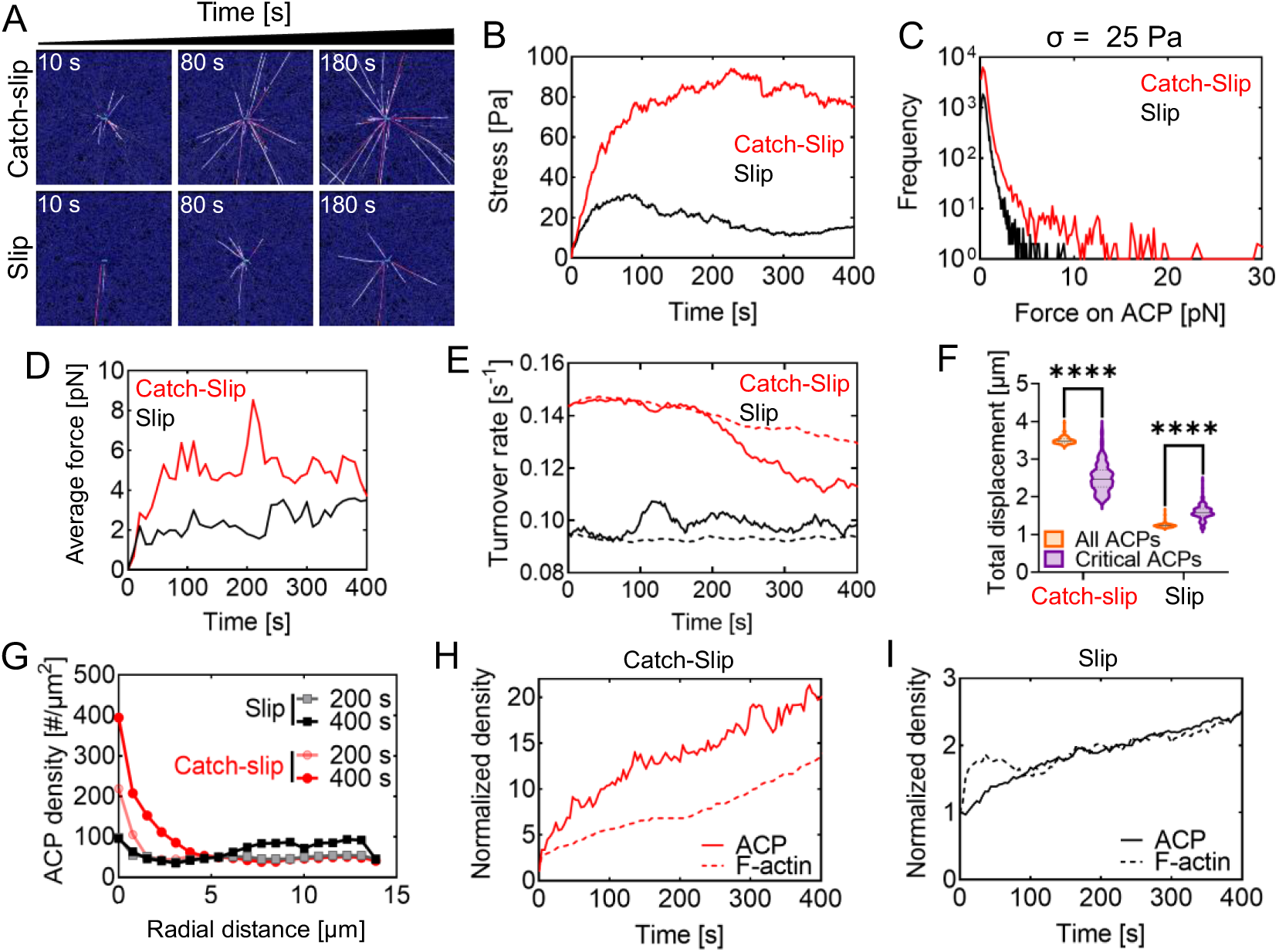
Catch-slip bonds enable molecular motors to generate higher contractile stress. **A-B,** A single immobile motor was placed at the center of the network to generate contractile forces in the presence of catch-slip-bond cross-linkers or slip-bond cross-linkers. The network with catch-slip-bond cross-linkers generated greater stress, and contractile forces propagated over longer distances, compared to those observed in the network with the slip-bond cross-linkers. **C,** Distribution of tensile forces acting on cross-linkers when stress was 25 Pa. **D,** Average tensile force acting on catch-slip-bond cross-linkers and slip-bond cross-linkers. **E-F,** The average turnover rate and total displacement of critical cross-linkers selected at the yield point (bearing top 10% of tensile forces). **G,** Density of cross-linkers as a function of radial distance at 200 s and 400 s. **H-I,** Normalized densities of filaments and cross-linkers near the motor. In the network with catch-slip-bond cross-linkers, cross-linker density increased faster than filament density, meaning that the redistribution of cross-linkers occurs earlier than filament aggregation induced by the motor. By contrast, the network with slip-bond cross-linkers showed filament aggregation took place earlier than cross-linker redistribution. Data are mean ± s.d., n=4. *n* values refer to individual simulation. Statistical analysis was performed using two-sided unpaired *t*-tests **(F)**. ns, not significant; \**P* < 0.05, \*\**P* < 0.01, \*\*\**P* < 0.001, \*\*\*\**P* < 0.0001.

We also tested whether catch-slip-bond cross-linkers and slip-bond cross-linkers show independent distinct behaviors in the same network. Simulations were repeated with both types of the cross-linkers (Fig. S8A). Stress generated by the network and the propagation distance of contractile forces were between those observed in two separate simulations run with each type of the cross-linker shown earlier (Fig. S8B). The catch-slip-bond cross-linkers sustained larger forces than the slip-bond cross-linkers (Fig. S8C). The average force and total displacement of the catch-slip-bond cross-linkers were significantly higher than those of the slip-bond cross-linkers (Figs. S8D, E). In addition, the catch-slip-bond cross-linkers were localized near the motor, whereas the slip-bond cross-linkers were more redistributed in locations far from the motor (Fig. S8F). These results demonstrate that both types of the cross-linkers exhibit their distinct behaviors independently. Note that it is possible to include both types of the cross-linkers in passive networks subjected to external strain, but we performed this test only for active networks because it is much easier to quantify their separation happening in the radial direction.

Overall, these simulations with the motor show that the force-dependent redistribution of cross-linkers takes place not only in the passive network but also in the active network. In addition, catch-slip-bond cross-linkers still enhance force-transmitting pathways in the active network by better connecting high-force bearing elements, thus increasing internal stress and force generated by the network. It is expected that variations in the three parameters defining the force-dependent lifetime of catch-slip bonds would lead to similar outcomes, which will be investigated in our future study. Unlike *γ* used for passive networks, there is no single parameter directly defining the rates of network deformation and force development on cross-linkers. However, it is likely that these rates are proportional to the walking rate of motors which is governed by mechanochemical rates in the crossbridge cycle [37–39]. Like our prediction in passive networks, it is expected that *τ*_0_ and *τ*_max_ need to be inversely proportional to the walking rate to result in similar outcomes.

## CONCLUSION

In this study, we explored how catch-slip bonds tune the material properties of cross-linked actin networks. Via extensive parametric studies, we demonstrated how the catch-slip bonds with different force dependence can vary stiffness and yield stress/strain over a wide range. In particular, we uncovered a distinct mechanism by which the catch-slip bonds enhance both yield stress and strain in networks with high connectivity. In this mechanism, the high-force bearing part keeps turning over, leading to relatively constant stiffness until the yield point with large stress and strain. Further, we demonstrated the catch-slip bonds govern the force generation process in active networks with molecular motors in a similar manner. Although our study was initially motivated by the actin cytoskeleton with a specific type of cross-linker, the mechanisms found in this study are general and applicable to other types of polymeric networks formed by catch-slip-bond cross-linkers that can be redistributed fast enough to adapt to force development. There have been several studies where artificial catch bonds were designed and tested using DNA or synthetic materials [40–42]. Based on these advances in molecular engineering, it would be feasible to design cross-linkers showing the nature of catch-slip bonds for many types of natural or synthetic polymers. For example, DNA-based cross-linkers have been already developed for actin filaments [40, 43], so transforming them to catch-slip bonds would not be hard. It could be also possible to change the profile of bond lifetime using different molecular structures. Then, our findings from simulations can be verified directly by such experimental systems. In addition, our study provides design principles for materials with desired mechano-adaptation properties.

## DATA AVAILABILITY STATEMENT

The data that support the findings of this study are available from the corresponding author upon reasonable request.

## FUNDING STATEMENT

We gratefully acknowledge the support from EMBRIO Institute, contract #2120200, a National Science Foundation (NSF) Biology Integration Institute.

## CONFLICT OF INTEREST DISCLOSURE

The authors declare no conflict of interest.

## AUTHOR CONTRIBUTIONS

M.F.R. and T.K. designed the project. M.F.R. performed simulations and analyzed data obtained from the simulations. T.K. supervised all computational studies. All the authors participated in writing the manuscript.

## Notes

### Competing Interest Statement

The authors have declared no competing interest.

