## Supporting Information for "Fine-Tuning of Material Properties via Catch Bonds"

### Simplification of actin filament and cross-linker

Actin filament, cross-linker, and motor are simplified using cylindrical segments (Fig. S1A). The actin filament is simplified into serially connected cylindrical segments whose length and diameter are 420 nm and 7 nm, respectively. Actin segments have polarity defined by barbed and pointed ends. Each cross-linker consists of two segments connected at its center point. Binding sites for the cross-linker segments are located on each actin segment every 7 nm. The number of the cross-linker segments that can simultaneously bind to one binding site is limited to two. The motor consists of a backbone structure with 64 motor arms; the backbone comprises 31 segments, and the endpoint of each backbone segment has two motor arms. The length of the motor backbone is 1.3 μm since each backbone segment is 42 nm long.

### Brownian dynamics via the Langevin equation

While the endpoints of motor backbone segments are fixed in space, segments for actin filament and cross-linker are displaced over time. The displacements of the endpoints of those mobile segments are governed by the Langevin equation with inertia neglected:

$$\mathbf{F}_i - \zeta_i \frac{\mathrm{d}\mathbf{r}_i}{\mathrm{d}t} + \mathbf{F}_i^{\mathrm{T}} = 0 \tag{S1}$$

where $\mathbf{F}_i$ is a net deterministic force, $\zeta_i$ is a drag coefficient, $\mathbf{r}_i$ is a position, $t$ is time, and $\mathbf{F}_i^{\mathrm{T}}$ is a stochastic force satisfying the fluctuation-dissipation theorem (1):

$$\left\langle \mathbf{F}_i^{\mathbf{T}}(t)\mathbf{F}_j^{\mathbf{T}}(t) \right\rangle = \frac{2k_{\mathbf{B}}T\zeta_i\delta_{ij}}{\Delta t}\boldsymbol{\delta} \tag{S2}$$

where $\delta_{ij}$ is the Kronecker delta, $\Delta t$ = 5.94×10$^{-5}$ s is a time step, $k_{\mathrm{B}}T$ is thermal energy, and $\boldsymbol{\delta}$ is a unit second-order tensor. $\zeta_i$ is calculated using the approximated equation for a cylindrical object (2):

$$\zeta_i = 3\pi\mu r_{c,i} \frac{3 + 2r_{0,i}/r_{\mathrm{c},i}}{5} \tag{S3}$$

where $\mu$ is the viscosity of a surrounding medium, and $r_{c,i}$ and $r_{0,i}$ are the diameter and length of a segment to which the endpoint belongs, respectively. In each time step, the positions of the endpoints of each segment are updated via the forward Euler integration scheme:

$$\mathbf{r}_i(t+\Delta t) = \mathbf{r}_i(t) + \frac{d\mathbf{r}_i}{dt}\Delta t = \mathbf{r}_i(t) + \frac{1}{\zeta_i}\left(\mathbf{F}_i + \mathbf{F}_i^{\mathrm{T}}\right)\Delta t \tag{S4}$$

**Deterministic forces for actin filament and cross-linker**

The deterministic force, $\mathbf{F}_i$, includes three kinds of forces: extensional, bending, and repulsive forces. The extensional and bending forces are calculated based on harmonic potentials:

$$U_{\mathrm{s}} = \frac{1}{2}\kappa_{\mathrm{s}}(r - r_0)^2 \tag{S5}$$

$$U_{\mathrm{b}} = \frac{1}{2}\kappa_{\mathrm{b}}\left(\theta - \theta_0\right)^2 \tag{S6}$$

where $\kappa_{\mathrm{s}}$ and $\kappa_{\mathrm{b}}$ are extensional and bending stiffnesses, respectively, $r$ is the length of a segment, $\theta$ is an angle formed by segments, and the subscript 0 represents an equilibrium value. The extensional ($\kappa_{\mathrm{s,F}}$) and bending ($\kappa_{\mathrm{b,F}}$) stiffnesses of filaments maintain the length of each actin segment near an equilibrium length ($r_{0,\mathrm{F}}$ = 420 nm) and an angle formed by adjacent actin segments near an equilibrium angle ($\theta_{0,\mathrm{F}}$ = 0°), respectively. The extensional ($\kappa_{\mathrm{s,X}}$) and bending ($\kappa_{\mathrm{b,X}}$) stiffnesses of cross-linkers maintain the length of each cross-linker segment near an equilibrium length ($r_{0,\mathrm{X}}$ = 23.5 nm) and an angle formed by two cross-linker segments near an equilibrium value ($\theta_{0,\mathrm{X}}$ = 0°), respectively. Forces exerted on the binding sites of the actin segments by the cross-linkers are distributed to two endpoints of the segments as explained in our previous work (3); a larger fraction of forces exerted by the cross-linkers are distributed to an endpoint closer to the binding site.

Repulsive forces acting between overlapping actin segments are calculated based on the following harmonic potential:

$$U_{\mathrm{r,F}} = \begin{cases} \frac{1}{2}\kappa_{\mathrm{r,F}}\left(r_{12,\mathrm{F}} - r_{\mathrm{c,F}}\right)^2 & \text{if} \quad r_{12,\mathrm{F}} < r_{\mathrm{c,F}} \\ 0 & \text{if} \quad r_{12,\mathrm{F}} \geq r_{\mathrm{c,F}} \end{cases} \tag{S7}$$

where $r_{12,\mathrm{F}}$ is a minimum distance between two neighboring actin segments, and $\kappa_{\mathrm{r,F}}$ is the strength of the repulsive force.

**Dynamics of actin filaments**

Due to the smaller dimension of the computational domain in z direction, the nucleation of filaments occurs in a random direction perpendicular to the z direction, followed by relatively fast polymerization. The nucleation of the filaments corresponds to the emergence of one actin segment occurring with a given nucleation rate constant ($k_{\mathrm{n,F}}$). The polymerization is represented by the addition of the actin segments to the barbed end of existing filaments with a given polymerization rate constant ($k_{\mathrm{p,F}}$). The depolymerization of actin filaments is not considered in this study.

**Dynamics of cross-linkers**

Cross-linkers exist in one of three states: monomeric, inactive, and active states. Monomeric cross-linkers are not bound to any filament, and they are considered implicitly by their local concentrations. The monomeric cross-linkers can bind to binding sites located on actin segments at a rate calculated using a binding rate constant, $k_{\mathrm{b,X}}$, and local monomer concentration. After binding, they transition to inactive cross-linkers bound to only one filament. The inactive cross-linkers can bind to binding sites located on another actin segment without any preference for a cross-linking angle at a rate determined by $k_{\mathrm{b,X}}$ if the following conditions are met. A distance between the binding site on the actin segment and the center point of the inactive cross-linker falls between 21.2 nm and 25.9 nm. After this second binding event, they transition to active cross-linkers forming functional cross-linking points between pairs of the filaments.

The active cross-linkers can also unbind from the filaments at a force-dependent rate described by Bell's law (4):

$$k_{\mathrm{u,X}} = \begin{cases} k_{\mathrm{u,X}}^{\mathrm{s0}} \exp\left( \dfrac{\lambda_{\mathrm{u,X}}^{\mathrm{s}} \left| \vec{F}_{\mathrm{s,X}} \right|}{k_{\mathrm{B}} T} \right) + k_{\mathrm{u,X}}^{\mathrm{c0}} \exp\left( \dfrac{-\lambda_{\mathrm{u,X}}^{\mathrm{c}} \left| \vec{F}_{\mathrm{s,X}} \right|}{k_{\mathrm{B}} T} \right) & \text{if } r \geq r_{0,\mathrm{X}} \\ k_{\mathrm{u,X}}^{\mathrm{s0}} + k_{\mathrm{u,X}}^{\mathrm{c0}} & \text{if } r < r_{0,\mathrm{X}} \end{cases} \tag{S8}$$

where $k_{\mathrm{u,X}}^{\mathrm{s0}}$ and $k_{\mathrm{u,X}}^{\mathrm{c0}}$ are zero-force unbinding rate constants for slip and catch bonds, respectively, and $\lambda_{\mathrm{u,X}}^{\mathrm{s}}$ and $\lambda_{\mathrm{u,X}}^{\mathrm{c}}$ are sensitivity to the magnitude of an applied spring force ($\left|\vec{F}_{\mathrm{s,X}}\right| = -\nabla U_{\mathrm{s,X}}$) for slip and catch bonds, respectively. In cases considering only slip bonds, $k_{\mathrm{u,X}}^{\mathrm{c0}}$ is set to zero. It is assumed that after the first unbinding event (i.e., active → inactive), the cross-linkers unbind quickly from the other filament and disappear (i.e., inactive → monomeric), and then they appear in a different location on a filament (i.e., monomeric → inactive) whose distance falls within 5 μm from the unbinding location. We implemented this event called "turnover" to account for the mobility of the cross-linkers. With higher mobility, the cross-linkers can be redistributed faster to adapt to a mechanical load that a network experiences. This turnover event occurs only once after the transition from the active cross-linkers to the inactive cross-linkers takes place. If the inactive cross-linkers emerging after the turnover event experience a transition to the monomeric state again via unbinding occurring at $k_{\mathrm{u,X}}^{\mathrm{s0}} + k_{\mathrm{u,X}}^{\mathrm{c0}}$ , they do not undergo the turnover event.

**Mechanics and dynamics of motors**

The extension of each motor arm attached to the endpoints of backbone segments is regulated by the two-spring model with the stiffnesses of transverse ($\kappa_{\mathrm{s,M1}}$) and longitudinal ($\kappa_{\mathrm{s,M2}}$) springs. The transverse spring regulates an equilibrium distance ($r_{0,\mathrm{M1}}$ = 13.5 nm) between the endpoint of the motor backbone and an actin segment where the arm of the motor is bound, and the longitudinal spring maintains a right angle between the motor arm and the actin segment ($r_{0,\mathrm{M2}}$ = 0 nm). Forces exerted on the actin segments by bound motors are also distributed to the barbed and pointed ends of the actin segments.

The motor is created by interconnecting 31 backbone segments at the center of the domain. The backbone is oriented in the y direction. Each motor arm represents the kinetics and force-velocity relationship of the ensemble of 8 myosin heads ($N_{\mathrm{h}}$ = 8). The motor arms located on the endpoints of the backbone segments bind to binding sites on actin segments at a constant rate, $40N_{\mathrm{h}}$ $\mathrm{s}^{-1}$. The walking ($k_{\mathrm{w,M}}$) and unbinding ($k_{\mathrm{u,M}}$) rates of the motor arms are determined by the parallel cluster model to account for the mechanochemical cycle of non-muscle myosin II (5, 6). The details of implementation and benchmarking of the parallel cluster model in our models were extensively described in our previous study (3). $k_{w,M}$ and $k_{u,M}$ in the model are lower with a larger applied load, based on an assumption that the motors exhibit the catch-bond behavior. The unloaded walking velocity and stall force of the motor arms are set to ~140 nm/s and ~6 pN, respectively.

**Network assembly**

At the beginning of each simulation, a cross-linked actin network is assembled via the dynamic events of actin filaments and cross-linkers in a three-dimensional rectangular domain (30×30×1 μm) with the periodic boundary condition in all directions (Fig. S1b). The slow nucleation and relatively fast polymerization of the filaments in the absence of depolymerization result in the formation of long filaments whose length shows the Gaussian-like distribution. The average length of the filaments in all simulations is ~12 μm, which is close to the typical average length of filaments in reconstituted networks (7, 8). The cross-linkers bind to pairs of filaments to form functional cross-linking points between the filaments. After forming the network, the periodic boundary condition is deactivated in the y direction, and the filaments crossing two boundaries normal to the y direction are severed and clamped to the boundaries. The filaments do not undergo further dynamic events.

**Bulk rheology simulations**

For the bulk rheology simulations, shear strain linearly increasing at the rate of $\dot{\gamma}$ = 0.001 $s^{-1}$ is applied to the +y boundary in the +x direction. Actin segments clamped to the +y boundary during the network assembly are displaced following the linearly increasing shear strain. The resultant stress generated by the network is calculated by summing the x-component of forces acting on the ends of the filaments clamped to the +y boundary and then dividing this sum by the area of the +y boundary. Due to the challenge of precisely controlling stress without a noise in simulations, the strain is chosen as an input parameter in the bulk rheology simulations. To identify yield points, we assessed the slope of the stress-strain curve at each time point to find when the slope starts deviating from its steady value. If the slope increases by >5% of the steady value, we consider that specific point the yield point.

**Motor contraction simulations**

For the motor contraction simulations, we place a single motor which resembles the myosin thick filament at the center of the network and let the motor generate internal contractile forces to the surrounding network by interacting with actin filaments. During the measurement, the backbone of the motor is fixed in space, whereas motor arms keep binding and walking on the actin filaments. Internal stress generated by the motor is measured as follows. First, the domain is evenly divided into 10 subdomains in the x direction. All chains of the filaments and cross-linkers that cross the 10 cross-sections between these subdomains are identified. Then, the x component of spring forces acting on those identified chains for each cross-section is summed, and the sum is divided by the area of the cross-section to calculate stress for each cross-section. This process is repeated in the y direction. The stress values calculated on the 20 cross-sections are averaged into a single stress value.

**Quantification of the number of cross-linkers per filament pair**

To understand the force-dependent distribution of cross-linkers on a network, we measure tensile forces acting on the active cross-linkers (bound to pairs of filaments) as a function of spring forces acting on filament pairs. Specifically, the spring forces acting on the two filaments are used as x and y coordinates in heat maps that show the level of the tensile forces acting on the active cross-linkers. 10×10 bins are used for the heat maps, meaning that the spring forces acting on the filaments belong one of the 10 bins. The size of bins is set in a non-uniform manner in order to have each bin contain 10% of the filaments. The bicubic interpolation is applied to create continuous, smooth color transitions on the heat maps.

**Analysis of the forces and turnover rate of cross-linkers**

To quantify forces acting on a subset of cross-linkers, the cross-linkers bearing the top 10% of tensile forces are identified at certain strain level. If this selection is made at the yield point, the selected cross-linkers are called critical cross-linkers. Once these cross-linkers are selected, their average spring force is traced across all strain levels, which is similar to the Lagrangian method (c.f., the Eulerian method) in terms of tracking the same entities over time. This allowed for monitoring the time evolution of forces developed on these specific cross-linkers as network deformation increases. The force distribution on these selected cross-linkers is then compared to the global average of forces acting on all cross-linkers in the network.

The turnover rate of the selected cross-linkers is also analyzed. The turnover rate is calculated by dividing the number of the turnover events of the cross-linkers occurring within a time window by the duration of the time window. The turnover rate is also compared with the global average calculated using all cross-linkers.

**Quantification of the densities of filaments and cross-linkers**

To quantify the densities of filaments and cross-linkers in the motor contraction simulations, we analyze their spatial distribution in the specific annular regions of the network. First, the network is divided into 19 concentric annuli (along xy plane) whose center is equal to the center of the network. For each annulus, the total numbers of the filaments and the cross-linkers are counted. The density in each annulus is then calculated by dividing the counted total numbers by the area of the annulus. Then, these density values are plotted as a function of a radial distance from the center of the motor (i.e., the network center).

**Supporting Figures**

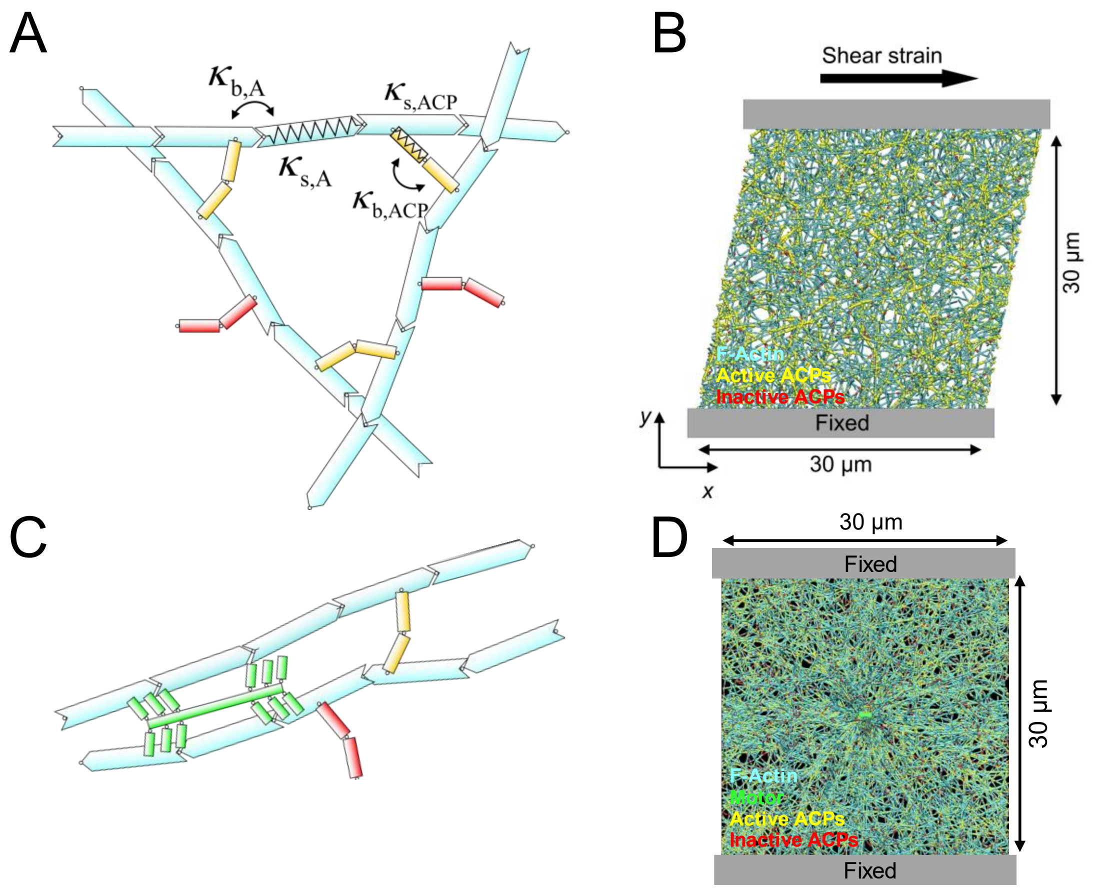


**Figure S1. Models for simulating passive and active actin networks. A,** Schematic showing a model for a passive actin network without motors. Actin filaments (cyan) are simplified into serially connected cylindrical segments. Cross-linkers are represented by two arm segments connected by elastic hinges. Cross-linkers bound to two filaments form functional cross-links (yellow), whereas those bound to only one filament are in an inactive state (red). The mechanics of these segments is governed by bending ($\kappa_b$) and extensional ($\kappa_s$) stiffnesses. The subscripts "F" and "X" on stiffnesses represent actin filaments and cross-linkers, respectively. **B,** Snapshot of the passive network (30 × 30 × 1 μm) subjected to linearly increasing shear strain for the bulk rheology measurement, where the -y boundary of the network is fixed, and the +y boundary is displaced in the +x direction**. C,** Schematic showing a model for an active actin network with a single motor (green) simplified into a backbone structure with motor arms. The length of the backbone is 1.3 μm, and the number of motor arms is 64. **D,** Snapshot of the active network (30 × 30 × 1 μm) with the motor located at the center of the network. Both +y and -y boundaries are fixed.

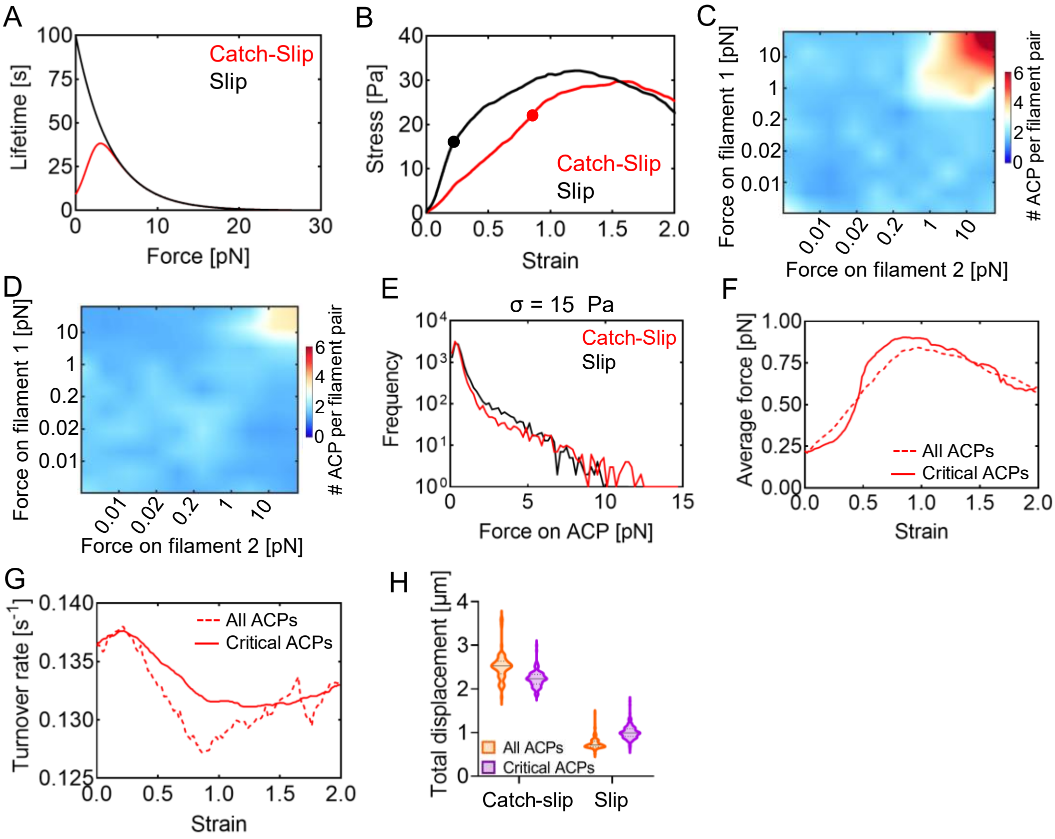


**Figure S2. Catch-slip-bond cross-linkers with different force dependence still strengthen actin networks. A,** Bond lifetime. The slip part of the lifetime of the catch-slip bond is identical to the lifetime of the slip bond. Thus, area under the lifetime curve of the catch-slip bond is smaller than the other. **B,** The stress-strain relationship of networks with the catch-slip-bond cross-linkers (red) and the slip-bond cross-linkers (black). Solid circles indicate the yield point in each network. **C-D,** The average number of cross-linkers bound to pairs of filaments bearing different tensile forces. **E,** The distribution of tensile forces on all cross-linkers when network stress was 15 Pa. **F-H,** A small group of cross-linkers, bearing the top 10% of tensile forces at the yield point, were selected and named critical cross-linkers. The behaviors of critical cross-linkers were compared with those of all cross-linkers. **F**, The average force acting on catch-slip-bond cross-linkers. **G,** The average turnover rate of the catch-slip-bond cross-linkers (red). **H,** The total displacement of cross-linkers.

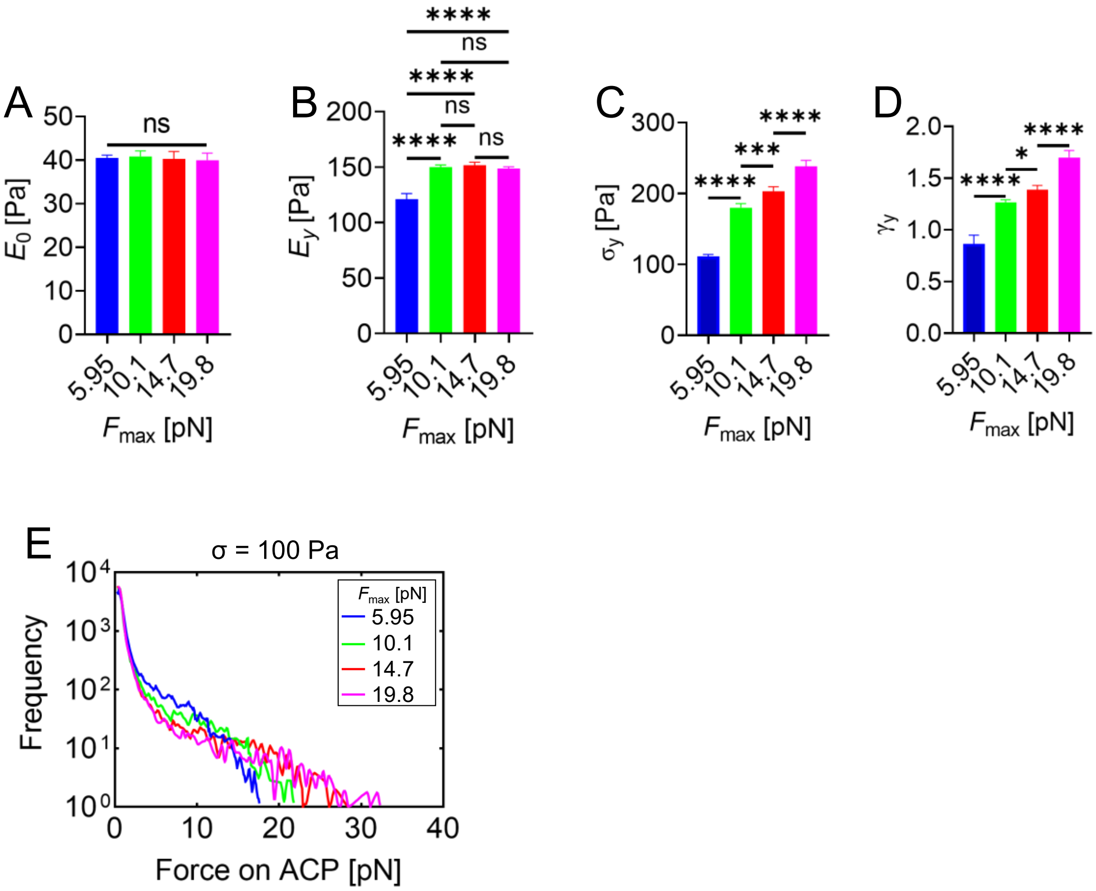


**Figure S3. Impact of varying $F_{max}$ on network stiffness and yield properties. A,** The stress-strain relationship at low stress and strain showed similar magnitudes and slopes across all conditions with varying Fmax, indicating comparable initial network stiffness ($E_0$). **B,** Network stiffness near the yield point ($E_y$) was consistent across conditions, except for the case with the lowest Fmax. **C,** Quantification of average yield stress for each condition, presented as mean ± s.d. (N = 4 independent samples per condition). **D,** Quantification of average yield strain for each condition, presented as mean ± s.d. (N = 4 independent samples per condition). **E,** Conditions with different maximal lifetime forces (Fmax) from Fig. 3, with measurements taken at a stress of 100 Pa. Data are mean ± s.d., n=4. *n* values refer to individual simulation. Statistical analysis was performed using one-way analysis of variance (ANOVA) followed by Tukey's multiple-comparison test **(A-D)**. ns, not significant; *$P < 0.05$, **$P < 0.01$, ***$P < 0.001$, ****$P < 0.0001$.

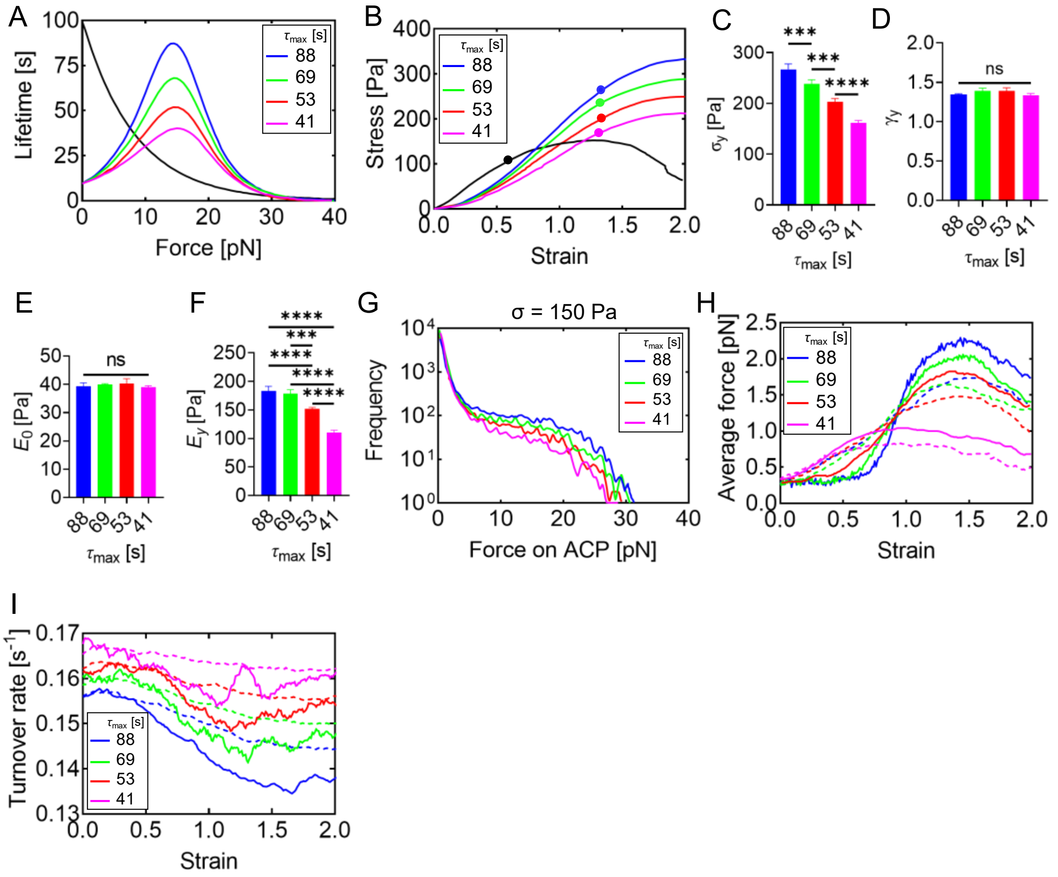


**Figure S4. Networks exhibit higher yield stress but similar yield strain if the maximal lifetime of catch-slip bonds is longer. A,** The lifetime of catch-slip bonds with different maximal lifetime ($\tau_{max}$). The black line shows the lifetime of the slip bond as a reference. **B,** The stress-strain relationship with different $\tau_{max}$. The black line shows the stress-strain relationship of the network with slip-bond cross-linkers. Solid circles indicate the yield point in each network. With longer $\tau_{max}$, network stiffness indicated by a tangent modulus (1/slope) was higher, whereas yield strain was similar in all cases. **C,** Quantification of average yield stress for each condition, presented as mean ± s.d. (N = 4 independent samples per condition). **D,** Quantification of average yield strain for each condition, presented as mean ± s.d. (N = 4 independent samples per condition). **E,** The stress-strain relationship at low stress and strain showed similar magnitudes and slopes across all conditions with varying $\tau_{max}$, indicating comparable initial network stiffness ($E_0$). **F,** Network stiffness near the yield point ($E_y$) was apparently larger with higher $\tau_{max}$. **G,** The impact of $\tau_{max}$ was analyzed by examining the tension distribution of filaments under 150 Pa stress. As $\tau_{max}$ increases, a greater number of catch-slip bonds withstand higher filament tension. Differences in frequency distribution are subtle but become significant under higher stress. **H,** The average force acting on all cross-linkers (dashed lines) and critical cross-linkers (solid lines) which withstand the top 10% of tensile forces at the yield point. **I,** The average turnover rate of all cross-linkers (dashed lines) and critical cross-linkers (solid lines). With longer $\tau_{max}$, deviations of the average force and turnover rate of the critical cross-linkers from the global averages were more substantial. Data are mean ± s.d., n=4. *n* values refer to individual simulation. Statistical analysis was performed using one-way analysis of variance (ANOVA) followed by Tukey's multiple-comparison test **(C-F)**. ns, not significant; $*P < 0.05$, $**P < 0.01$, $***P < 0.001$, $****P < 0.0001$.

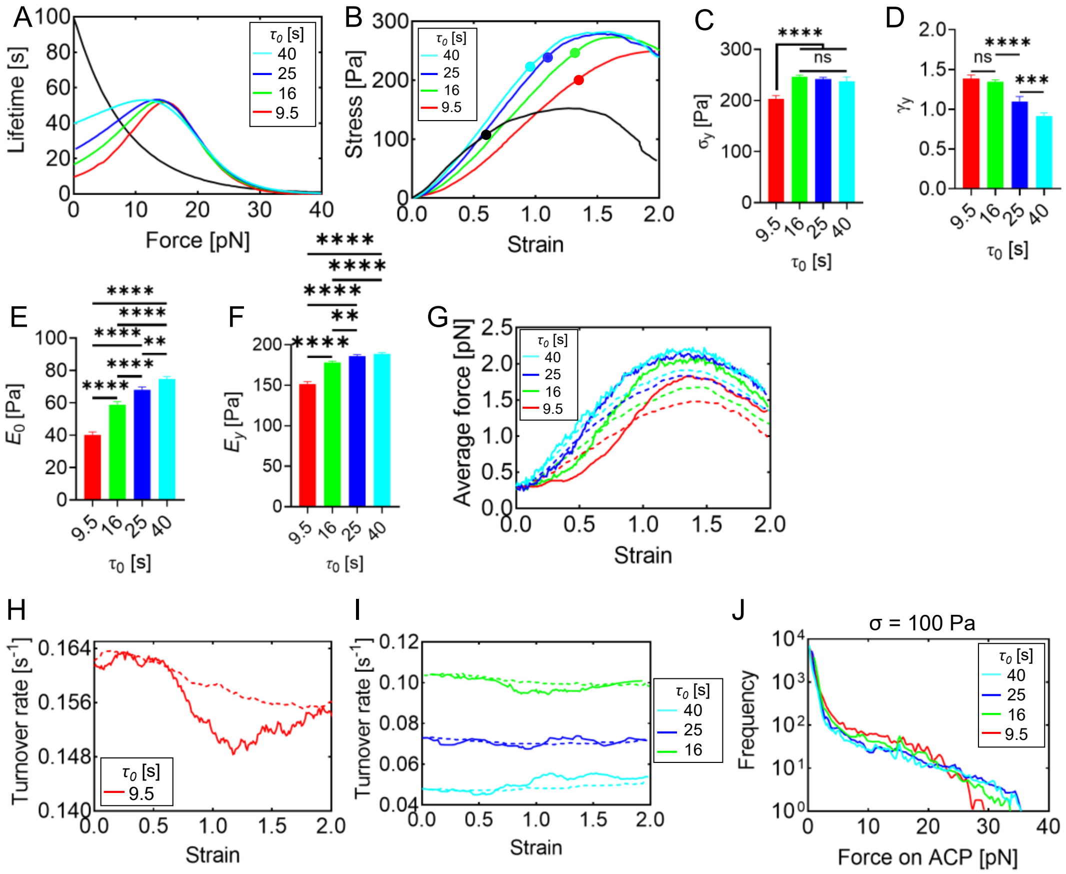


**Figure S5. Comparison of catch-slip bonds with different lifetime with zero force ($\tau_0$). A,** The lifetime of catch-slip bonds with different lifetime with zero force ($\tau_0$). **B,** The stress-strain relationship at different $\tau_0$ values. Solid circles represent the yield point for each network. As $\tau_0$ increases, the network stiffness (indicated by the tangent modulus, 1/slope) becomes higher, while the yield strain decreases. **C,** Quantification of the average yield stress for each condition, reported as the mean ± s.d. (N = 4 independent samples per condition). **D,** Quantification of the average yield strain for each condition, reported as the mean ± s.d. (N = 4 independent samples per condition). **E,** A notable difference in $E_0$ was observed, with higher $\tau_0$ corresponding to increased $E_0$. **F,** Network stiffness near the yield point ($E_y$) appeared larger at higher $\tau_0$. **G,** The average force on all cross-linkers (dashed lines) and the critical cross-linkers (solid lines) that bear the top 10% of tensile forces at the yield point. **H-I,** The average turnover rate of all cross-linkers (dashed lines) and critical cross-linkers (solid lines). As $\tau_0$ increases, the turnover rate decreases, with minimal deviation from the global average. **J,** The impact of $\tau_0$ was analyzed by examining the tension distribution of filaments under 100 Pa stress. As $\tau_0$ increases, a greater number of catch-slip bonds withstand higher filament tension.

Data are mean ± s.d., n=4. *n* values refer to individual simulation. Statistical analysis was performed using one-way analysis of variance (ANOVA) followed by Tukey's multiple-comparison test (C-F). ns, not significant; $*P < 0.05$, $**P < 0.01$, $***P < 0.001$, $****P < 0.0001$.

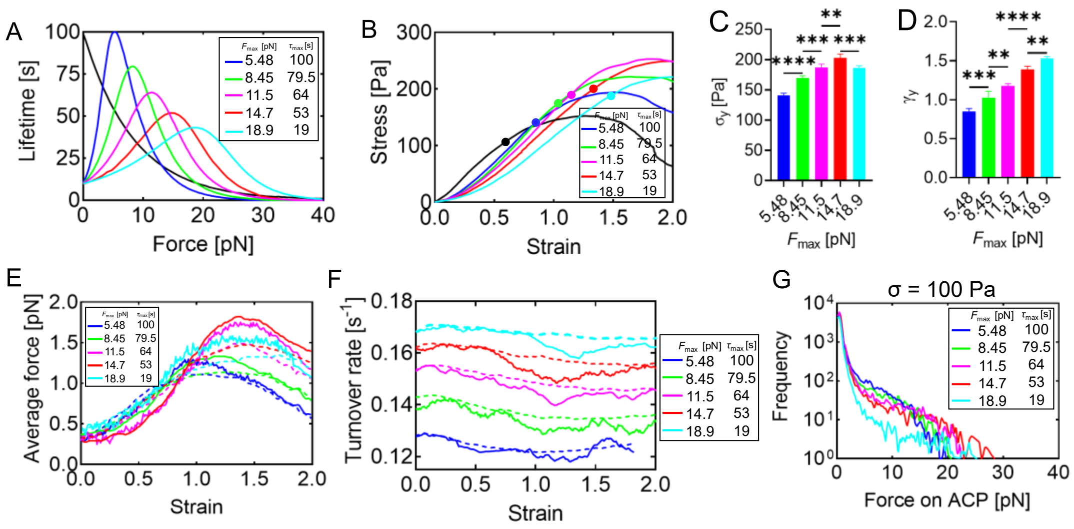


**Figure S6. Comparison of catch-slip bonds with different force dependence and the same area under the lifetime curve. A,** The lifetime of catch-slip bonds with different maximal lifetime ($\tau_{max}$) and different force for maximal lifetime ($F_{max}$). Area under all the lifetime curves is identical. The black line shows the lifetime of the slip bond as a reference. **B,** The stress-strain relationship. The black line shows the stress-strain relationship of the network with slip-bond cross-linkers. Solid circles indicate the yield point in each network. The case with intermediate values of $\tau_{max}$ and $F_{max}$ exhibited the highest yield stress and strain. **C,** Quantification of the average yield stress for each condition, reported as the mean ± s.d. (N = 4 independent samples per condition). **D,** Quantification of the average yield strain for each condition, reported as the mean ± s.d. (N = 4 independent samples per condition). **E,** The average force acting on all cross-linkers (dashed lines) and critical cross-linkers (solid lines) which withstand the top 10% of tensile forces at the yield point. **F,** The average turnover rate of all cross-linkers (dashed lines) and critical cross-linkers (solid lines). G, The distribution of tensile forces acting on cross-linkers when stress was 100 Pa. Data are mean ± s.d., n=4. *n* values refer to individual simulation. Statistical analysis was performed using one-way analysis of variance (ANOVA) followed by Tukey's multiple-comparison test **(C-D)**. ns, not significant; *$P < 0.05$, **$P < 0.01$, ***$P < 0.001$, ****$P < 0.0001$.

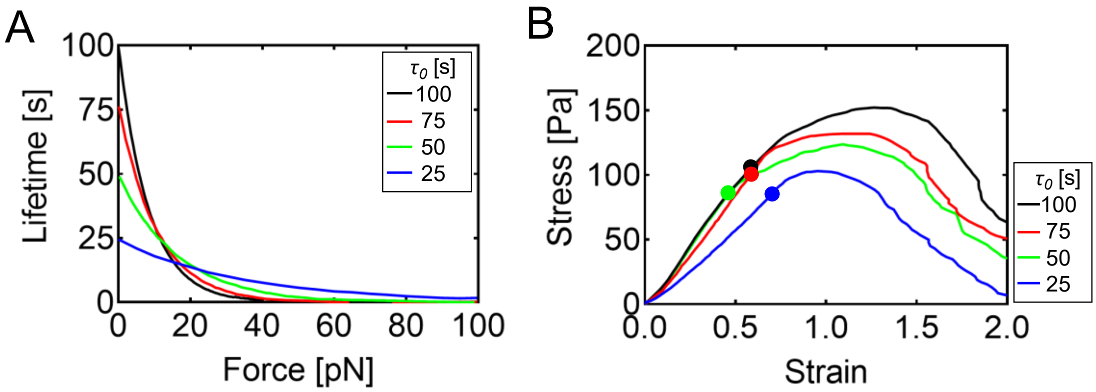


**Figure S7. Comparison of slip bonds with different lifetime with zero force ($\tau_0$) and the same area under the lifetime curve. A,** The lifetime of slip bonds with different lifetime with zero force ($\tau_0$). Area under all the lifetime curves is identical. **B,** The stress-strain relationship. The black line shows the stress-strain relationship of the network with slip-bond cross-linkers. Solid circles indicate the yield point in each network. As $\tau_0$ decreases, the yield point shifts to lower values relative to the reference condition. For the lowest $\tau_0$, we observed a decrease in Young's modulus and the yield strain was not changed significantly.

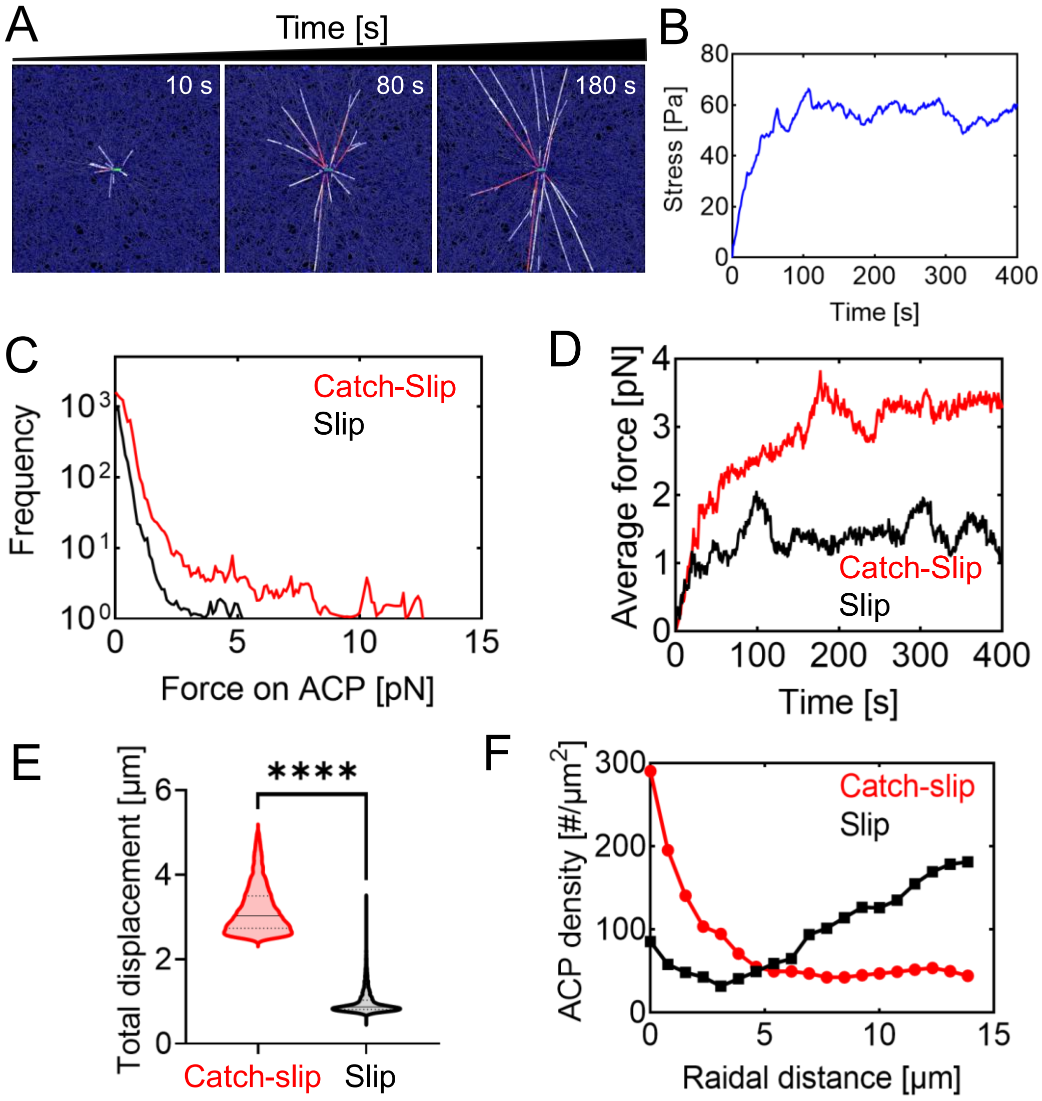


**Figure S8. Different behaviors of catch-slip bonds and slip-bonds in the same network. A-B,** A single immobile motor was placed at the center of the network to generate contractile forces with both catch-slip-bond cross-linkers and slip-bond cross-linkers. Stress generated by the network and the propagation distance of contractile forces are between two cases shown in Fig. 6. **C,** Distribution of tensile forces acting on cross-linkers. **D,** Average tensile force acting on cross-linkers. **E,** The total displacement of cross-linkers. **F,** Densities of cross-linkers as a function of radial distance from the motor. Data are mean ± s.d., n=4. *n* values refer to individual simulation. Statistical analysis was performed using two-sided unpaired *t*-tests **(E)**. ns, not significant; *$P < 0.05$, **$P < 0.01$, ***$P < 0.001$, ****$P < 0.0001$.

**Table S1.** List of parameters used in the model. References are provided for some of the parameters. "*" indicates the reference values of parameters.

| Symbol | Definition | Value |
|---|---|---|
| $r_{0,\mathrm{F}}$ | Length of an actin segment | $4.2\times10^{-7}$ [m] |
| $r_{c,\mathrm{F}}$ | Diameter of an actin segment | $7.0\times10^{-9}$ [m] (9) |
| $\theta_{0,\mathrm{F}}$ | Bending angle formed by adjacent actin segments | 0 [rad] |
| $\kappa_{s,\mathrm{F}}$ | Extensional stiffness of actin filaments | $1\times10^{-3}$ [N/m] |
| $\kappa_{b,\mathrm{F}}$ | Bending stiffness of actin filaments | $8.8\times10^{-20}$ [N·m] (10) |
| $r_{0,\mathrm{X}}$ | Length of a cross-linker arm | $2.35\times10^{-8}$ [m] (11) |
| $r_{c,\mathrm{X}}$ | Diameter of a cross-linker arm | $1.0\times10^{-8}$ [m] |
| $\theta_{0,\mathrm{X}}$ | Bending angle formed by two cross-linker arms | 0 [rad] |
| $\kappa_{s,\mathrm{X}}$ | Extensional stiffness of cross-linkers | $1.0\times10^{-3}$ [N/m] |
| $\kappa_{b,\mathrm{X}}$ | Bending stiffness of cross-linkers | 0 [N·m] |
| $r_{0,\mathrm{M1}}$ | Length of the transverse spring for a motor arm | $1.35\times10^{-8}$ [m] |
| $r_{0,\mathrm{M2}}$ | Length of the longitudinal spring for a motor arm | 0 [m] |
| $r_{c,\mathrm{M}}$ | Diameter of a motor arm | $1.0\times10^{-8}$ [m] |
| $N_\mathrm{h}$ | Number of heads represented by a motor arm | 8 |
| $N_\mathrm{a}$ | Number of arms on the motor | 64 |
| $\kappa_{s,\mathrm{M1}}$ | Extensional stiffness for the transverse spring for a motor arm | $1.0\times10^{-3}$ [N/m] |
| $\kappa_{s,\mathrm{M2}}$ | Extensional stiffness for the longitudinal spring for a motor arm | $1.0\times10^{-3}$ [N/m] |
| $k_{n,\mathrm{F}}$ | Nucleation rate of actin | 0.001 [$\mu M^{-1}s^{-1}$] |
| $k_{p,\mathrm{F}}$ | Polymerization rate of actin at the barbed end | 60000 [$\mu M^{-1}s^{-1}$] |
| $k_{b,\mathrm{X}}$ | Binding rate of cross-linkers | 100 [$\mu M^{-1}s^{-1}$] |
| $k_{u,\mathrm{X}}^{s0}$ | Zero-force unbinding rate constant of cross-linkers for slip | 0.0002 [$s^{-1}$] * |
| $k_{u,\mathrm{X}}^{c0}$ | Zero-force unbinding rate constant of cross-linkers for catch | 0.1 [$s^{-1}$] * |
| $\lambda_{u,\mathrm{X}}^{s}$ | Sensitivity of cross-linker unbinding to applied force for slip | $1.0\times10^{-9}$ [m] * |
| $\lambda_{u,\mathrm{X}}^{c}$ | Sensitivity of cross-linker unbinding to applied force for catch | $0.6\times10^{-9}$ [m] * |
| $\kappa_{r,\mathrm{F}}$ | Strength of repulsive force between actin filaments | $1.69\times10^{-3}$ [N/m] |
| $\Delta t$ | Time step | $5.94\times10^{-5}$ [s] |
| $M$ | Viscosity of medium | $8.6\times10^{-1}$ [kg/m·s] |
| $k_\mathrm{B}T$ | Thermal energy | $4.142\times10^{-21}$ [J] |
| $C_\mathrm{F}$ | Filament concentration | 5 [μM] |
| $R_\mathrm{X}$ | Ratio of cross-linker concentration to filament concentration | 0.01 |
| $\langle L_\mathrm{F} \rangle$ | Average filament length | ~12 [μm] |
| $\dot{\gamma}$ | Shear strain rate | 0.001 [$s^{-1}$] |
